# Wampee chromosome-level reference genome elucidates fruit sugar-acid metabolism

**DOI:** 10.1101/2024.04.16.589530

**Authors:** Huiqiong Chen, Jingxuan Wang, Xiangfeng Wang, Cheng Peng, Xiaoxiao Chang, Zhe Chen, Bowen Yang, Xinrui Wang, Jishui Qiu, Li Guo, Yusheng Lu

## Abstract

Wampee (*Clausena lansium*) is an economically significant subtropical fruit tree widely cultivated in Southern China. High-quality genomic resources are unavailable, but they are essential for functional genomics and germplasm enhancement of wampee. Here, we provide a chromosome-level genome sequence for the wampee cultivar JinFeng and a population genomic analysis of 266 accessions. The 297.1 Mb wampee genome, containing nine chromosomes with a scaffold N50 of 29.2 Mb and encoding 23,468 protein-coding genes, showed a significant improvement over the previous version. We dissected the wampee population structure and genetic differentiation in China using population genomic analysis, which detected 110 and 671 genes under a selective sweep associated with sour and sweet wampee evolution in domesticated clones, respectively. Homozygous non-synonymous single nucleotide polymorphisms are likely associated with fruit flavor differentiation. A genome-wide association study identified 220 remarkable marker-trait associations for total acid content, harboring 289 genes encoding transcription factors, transporters, and enzymes involved in sugar and acid metabolism, which are potentially useful for sour and sweet taste development in wampee fruit. Furthermore, the ethylene response factor family gene *ClERF061* and the SWEET family gene *ClSWEET7* were identified. Linkage assessment between the relative expression levels of *ClERF061* or *ClSWEET7* and the total acid/total sugar contents implied their potential involvement in sugar-acid metabolism in wampee fruits. High-quality genome resources are valuable for expediting wampee research and genome-assisted breeding.

## Introduction

Wampee (*Clausena lansium*), a fruit tree from the Rutaceae family, is typically grown in Southern China, particularly in the Guangdong, Fujian, Hainan, and Guangxi provinces, and sometimes found in Sri Lanka, Australia, India, America, and South Asia ^1,2^. The *Clausena* genus comprises approximately 30 species scattered across tropics and subtropics, with around 10 species and two varieties indigenous to Southern China ^3,4^. China has a rich history of cultivating and utilizing wampee for over 1500 years, first recorded in the ancient Chinese agricultural publication ‘*Qimin Yaoshu*’ in 533-544 AD), resulting in abundant germplasm resources. Wampee has been used as food and medicine in traditional Chinese medicine to cure ailments including abdominal pain, malaria, colds, dermatopathies, and snakebites ^5,6^. In the past few years, modern medical studies have indicated that natural products extracted from wampees, including flavonoids, coumarins, and alkaloids, show pharmaceutical properties as antioxidants, tumor inhibitors, and potential prevention of age-related cognitive decline ^7^. Apart from its medicinal and culinary attributes, wampee is also appreciated for its distinctive flavor, which has gained popularity among consumers and has economic benefits for local fruit growers.

Traditionally, wampees have been cultivated scattered throughout homes, resulting in diverse germplasm resources. However, the population genetic structure of wampees resulting from both inbreeding and crossbreeding remains poorly elucidated. Previous analyses employing markers, including random amplified polymorphic DNA or inter-simple sequence repeats, have suggested a close relationship between wampee varieties in Guangdong and Guangxi provinces. However, these markers have not provided a conclusive classification of the varieties owing to variations in marker selection, limited number of varieties examined, and differences in classification methods. Recently, a draft genome assembly of wampees was reported; however, it was highly fragmented, consisting of 45,747 contigs, with a contig N50 of 39 kb (Fan et al., 2021). To elucidate the genetic diversity and population structure of wampee varieties across China, it is imperative to obtain a high-quality reference genome sequence and comprehensively evaluate the genomic background. This information is vital for wampee conservation and breeding.

Fruit quality can be categorized into external and internal qualities. External quality primarily involves size, weight, color, and smoothness. Internal quality includes biochemical properties that reflect the nutritional value of fruits such as proteins, fats, vitamins, and minerals. They also include substances that reflect the flavor of the fruit, such as sugars, organic acids, tannins, and essential oils. Internal quality is an important indicator of the commercial quality of fruits, which primarily depends on the sugar-acid composition and sugar-acid ratio. The total sugar content of fruits varies little across different wampee varieties, but the organic acid content differs greatly, making it a determining factor affecting the sugar-acid ratio and flavor of the wampee fruit. Based on organic acid content, wampees can be divided into two subgroups with distinct taste profiles: extremely sweet and extremely sour. This extreme sweet-sour characteristic makes it an interesting choice for studying fruit sugar acid metabolism and sugar-acid ratio regulation.

In this study, we report a chromosome-level genome assembly for wampee by generating Illumina paired-end (PE), PacBio single-molecule real-time (SMRT), and high-throughput chromatin conformation capture (Hi-C) sequencing data from the ’JinFeng’ cultivar. Population sequencing analysis of 266 wampee accessions uncovered the genetic heterogeneity of the wampee population, demonstrating hybridization and cross-pollination among cultivars from major wampee-growing regions. Furthermore, selective sweep and genome-wide association study (GWAS) analyses determined multiple genes related to total acid content in wampee fruit, providing insights into the genome architecture and adaptation processes in this species. Genome assembly, annotation, and genetic variations are beneficial resources for functional genomic research and targeted wampee breeding aimed at enhancing fruit quality and investigating its potential medicinal properties.

## Results

### Chromosome-level genome assembly and characteristic analyses

To create a chromosome-level reference genome for *C. lansium* (2n = 18) (**Figure S1**), 141 Gb PacBio SMRT reads (∼491.29×coverage), 89 Gb (∼310.10×coverage) Illumina PE reads, and 57.6 Gb (∼127× coverage) Hi-C Illumina read pairs (**Table S1**) were produced from the *C. lansium* cultivar ’JinFeng’ altogether (**Figure 1A**). The cultivar has an approximated genome size of 284.15 Mb and a heterozygosity rate of 0.53% based on k-mer frequency analysis (**Figure S2**). The genome was assembled from PacBio SMRT reads using Falcon v0.7 ^8^, followed by contig polishing with Nextpolish v1.1.0 ^9^ using Illumina PE reads, resulting in a preliminary genome assembly of 297.07 Mb (**Table 1**). Hi-C PE reads were used to map contigs onto chromosomes using LACHESIS (v201701) ^10^. The ultimate genome assembly of ’JinFeng’ was 297.07 Mb in size, with 93.83% of the assembled sequences mapped onto nine pseudochromosomes (**Figure 1B**) with contig and scaffold N50 of 10.2 Mb and 29.2 Mb, separately (**Table 1**). This demonstrates a notable advancement beyond the previously announced assembly with a contig N50 of 0.039 Mb ^11^. The genome integrity was evaluated applying the Benchmarking Universal Single Copy Orthologs (BUSCO) database v5.3.2, revealing a 98.0% completeness for ’JinFeng’ (**Table 1, Table S2**). A total of 23,468 protein-coding genes and 4,177 non-coding RNAs (**Table 1**) were predicted in the ’JinFeng’ genome using EVidenceModeler, integrating information from *ab initio* prediction, protein alignments, and multi-tissue (root, seed, stem, leaf, flower, and fruit) transcriptome sequencing (**Table S3**). The GC content of ’JinFeng’ was 33.52% with a density of 79 genes per Mb (**Table 1**). Approximately 95.50% of the genes were labeled using the non-redundant protein sequence database, and 74.20% were labeled using Kyoto Encyclopedia of Genes and Genomes (KEGG) terms (**Table S4**). Repeat annotations showed that 46.48% of the ’JinFeng’ genome included repetitive elements, containing 33.32% long terminal repeat retrotransposons and 2.25% DNA transposons (**Table S5**).

**Figure 1.**
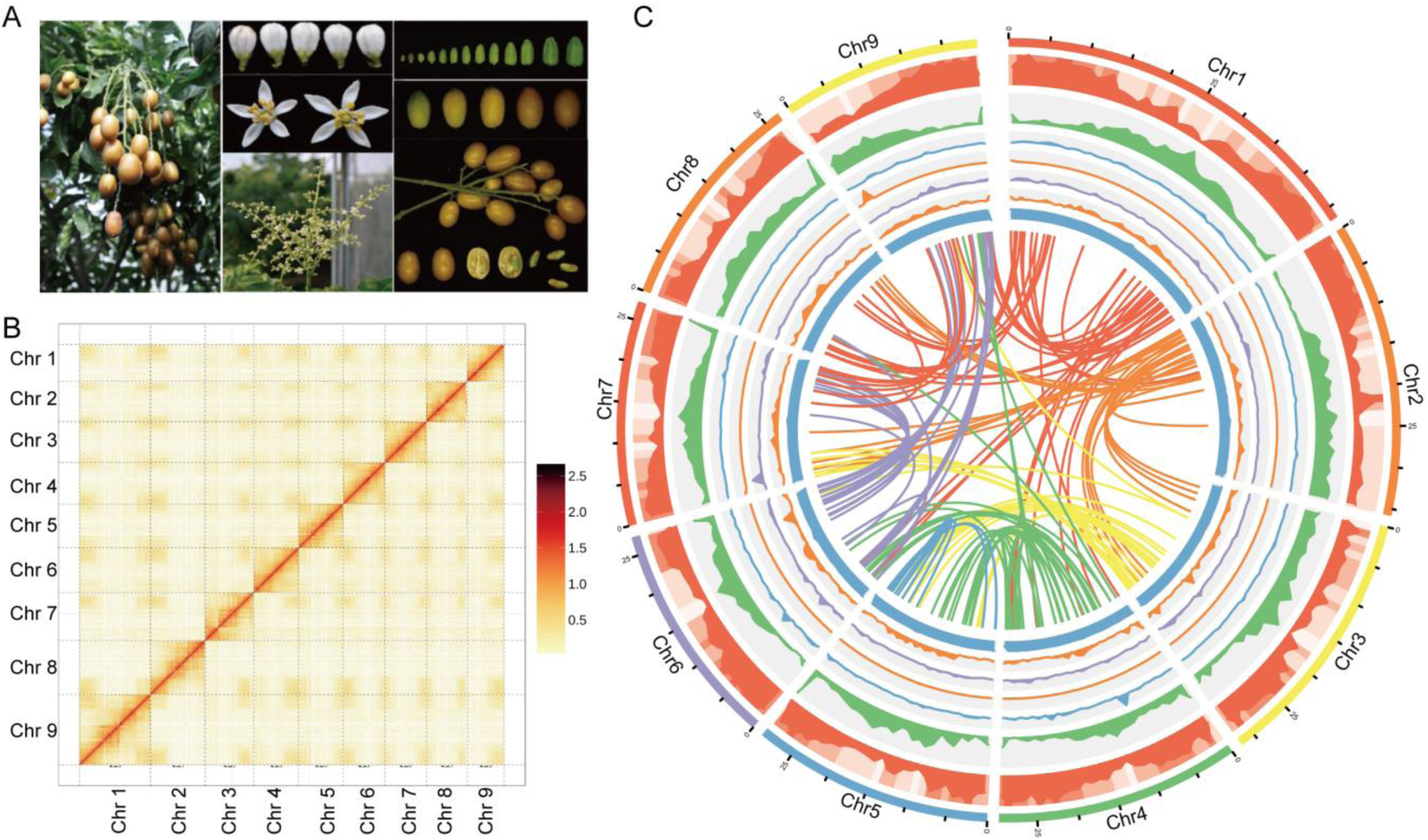
Chromosome-level assembly of wampee [*Clausena lansium*. (Lour.) Skeels] genome. (A) Photographs of wampee flowers and fruits at various developmental stages. (B) Heat map showing the density of the high-throughput chromosome conformation capture (Hi-C) interactions between chromosomes in *C. lansium*. (C) Overview of the wampee genome with the following features depicted from the exterior to the interior: chromosomes, gene density, transposable elements, distribution of GC content, distribution of miRNA density, distribution of tRNA density, distribution of snRNA density, distribution of rRNA density, paralogous relationships between wampee chromosomes.

**Table 1.** Data for *Clausena lansium* ’JinFeng’ genome assembly and annotations.

|  |  | 'JinFeng' | Fan *et al.*,<br>2021 |
| --- | --- | --- | --- |
| Contigs | Total Number | 163 | 45,747 |
|  | Assembly size | 297.1 Mb | NA |
|  | Contig N50 | 10.2 Mb | 39.37 Kb |
|  | Contig N90 | 1.2 Mb | 5.403 Kb |
|  | Largest Contig | 19.3 Mb | 303.82 Kb |
| Scaffolds | Total Number | 119 | 38,459 |
|  | Assembly size | 297.1 Mb | 310.51 Mb |
|  | Scaffold N50 | 29.2 Mb | 2.24 Mb |
|  | Scaffold N90 | 39 Mb | 6.779 Kb |
|  | BUSCO (%) | 98 | 94.7 |
| Annotation | Largest scaffold | 46.5 Mb | 15.9 Mb |
|  | Number of genes | 23,468 | 32,793 |
|  | Repeat content (%) | 46.48 | 39.08 |
|  | Number of cRNAs | 4,177 | 2,292 |
|  | GC content (%) | 33.5 | 37.44 |

### Comparative genomic and evolutionary analysis

Phylogenomic analysis of *C. lansium* and 11 angiosperm plants based on single-copy orthologs confirmed a close relationship between *C*. *lansium* and *Atalantia buxifolia*, *Poncirus trifoliata*, *Citrus sinensis*, and *Fortunella hindsii*. Using a molecular clock, we determined that *C. lansium* diverged from the ancestor of *F. hindsii* around 21.6 million years ago (Mya) and from the ancestor of *Dimocarpus longan* (longan) approximately 72.9 Mya (**Figure 2A**). Ortholog clustering analysis using OrthoFinder revealed 27,506 gene families, 6,134 of which were mutual among all 12 species, and 465 were single-copy gene families (5,580 genes) (**Figure 2B**). Specifically, we recognized 381 orthogroups unique to *C. lansium* compared with *C. sinensis* (sweet orange), *A. buxifolia* (boxthorn), *P. trifoliata* (trifoliate orange), and *F. hindsii* (Hong Kong kumquat) (**Figure 2C**). The expansion or contraction of gene families can offer valuable insights into the evolutionary forces that influence plant genomes and play significant roles in plant diversification. Gene family evolutionary analysis using CAFÉ identified 900 expanded and 1,512 contracted gene families (*P* ≤ 0.05) (**Table S6)**. Expanded gene families exhibited a significant enrichment of Gene Ontology terms associated with endopeptidase activity, regulation of translational initiation, nucleic acid transport, adenylyl sulfate kinase, and oxidoreductase activity. The expanded gene families were considerably enriched with KEGG terms for alkaloid biosynthesis, such as tropane, piperidine, isoquinoline, pyridine, monobactam biosynthesis, and tyrosine and phenylalanine metabolism (**Figure 2C; Table S7 and S8**).

**Figure 2.**
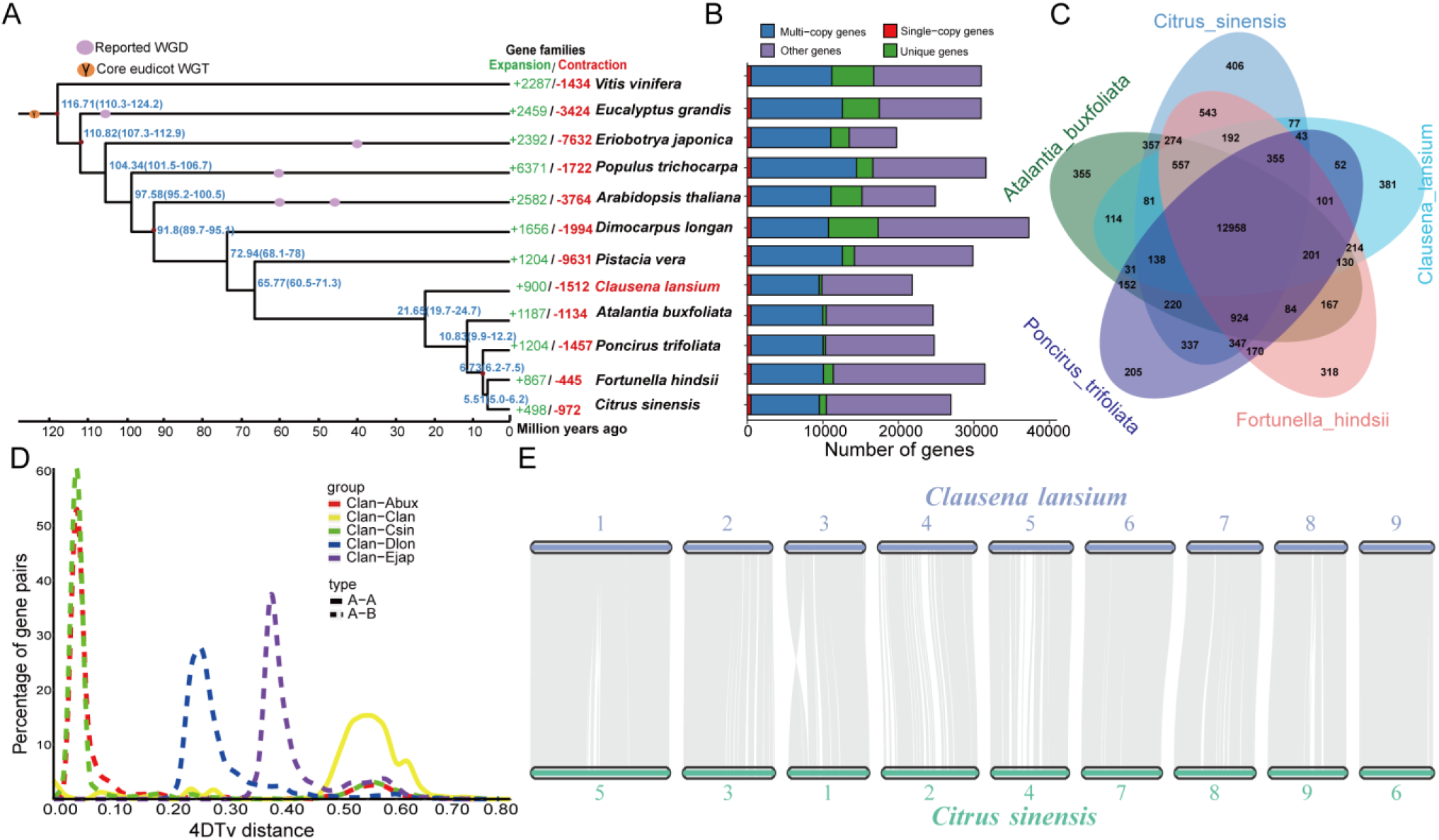
Evolution of *Clausena lansium* genome. (A) Phylogeny, estimated divergence time, and gene family expansions/contractions for 12 related species. Gene family expansion (green) and contraction (red) are illustrated. The red circles on certain nodes represent fossil calibration points acquired from the TIMETREE website (http://www.timetree.org/). Estimated divergence times (million years ago) are marked in blue at intermediate nodes, based on maximum likelihood analysis, with numbers in parentheses denoting 95% confidence intervals for the divergence time between different clades. Whole-genome duplication (WGD) and whole-genome triplication (WGT) events are denoted via colored dots. (B) Comparison of genes among 12 related species. (C) The shared and unique gene families of five species are illustrated using a Venn diagram. The number of genes in each family is denoted by the number in the corresponding color-coded area. (D) Four-fold degenerate synonymous sites of third codons (4DTv) analysis in *C. lansium* and related species. (E) The genomes of *Citrus sinensis* share syntenic blocks with *C. lansium*, with paired genes linked via grey lines.

Whole-genome duplication (WGD) is widespread in angiosperms and plays a key role in plant evolution. To analyze the evolutionary history and WGD events in the wampee genome, we calculated the 4-fold degenerate synonymous sites of third codons (4DTv) for the orthologs of *C. lansium* with *C. sinensis*, *Eriobotrya japonica* (loquat), *A. buxifolia*, and *D. longan,* which exhibited 4DTv distance peaks at 0.05, 0.05, 0.25, and 0.4, respectively. *C. lansium* diverged from *E. japonica*, *D. longan*, Rutaceae species *A. buxifolia*, and *C. sinensis* at around 104, 72.9 and 21.6 Mya, respectively, based on the MCMCtree (**Figure 2A**). Whole-genome triplication (γ), indicated by the 4DTv value, peaked at around 0.55 and occurred in the progenitor species of *C. lansium* (core dicotyledon) without additional WGDs afterwards. Synteny analysis also revealed a 1:1 collinearity between *C. lansium* and *C. sinensis*, as well as between *C. lansium* and *Vitis vinifera* (winegrape), confirming a lack of post-γ WGD in both genomes. Wampee and sweet orange exhibited very good collinearity, possibly because of their relatively recent divergence (**Figure 2E and S3**). Additionally, 1:2 collinearity was observed between *C. lansium* and *E. japonica*, confirming a recent WGD event in *E. japonica* (**Figure 2D and 2E**), as reported previously ^12^.

### Genetic diversity and population structure of wampee germplasm

To investigate the population structure and genetic diversity, we sequenced (∼40x coverage) a total of 266 wampee accessions, including 172 landraces and 94 cultivars collected from major wampee production areas across China, including 188 from Guangdong Province, 18 from Hainan, 1 from Yunnan, 38 from Guangxi, and 21 from Fujian (**Figure 3A, Table S9**). A total of 78,266,957 single-nucleotide polymorphisms (SNPs) were identified, 1,040,809 of which were high-quality SNPs (minor allele frequency > 5%) (**Figure S3 to S5 Table S10 to S12**). Phylogenomic and principal component analyses (PCA) using these high-quality SNPs revealed the population structure and genetic diversity within these wampee cultivars, where cultivated and landrace varieties were grouped separately overall, with most sweet varieties clustered together within the cultivated varieties. In contrast, varieties from various geographic regions lacked a clear separation, suggesting a complex history of wampee migration and a genetic mixture from different provinces (**Figure 3B**).

**Figure 3.**
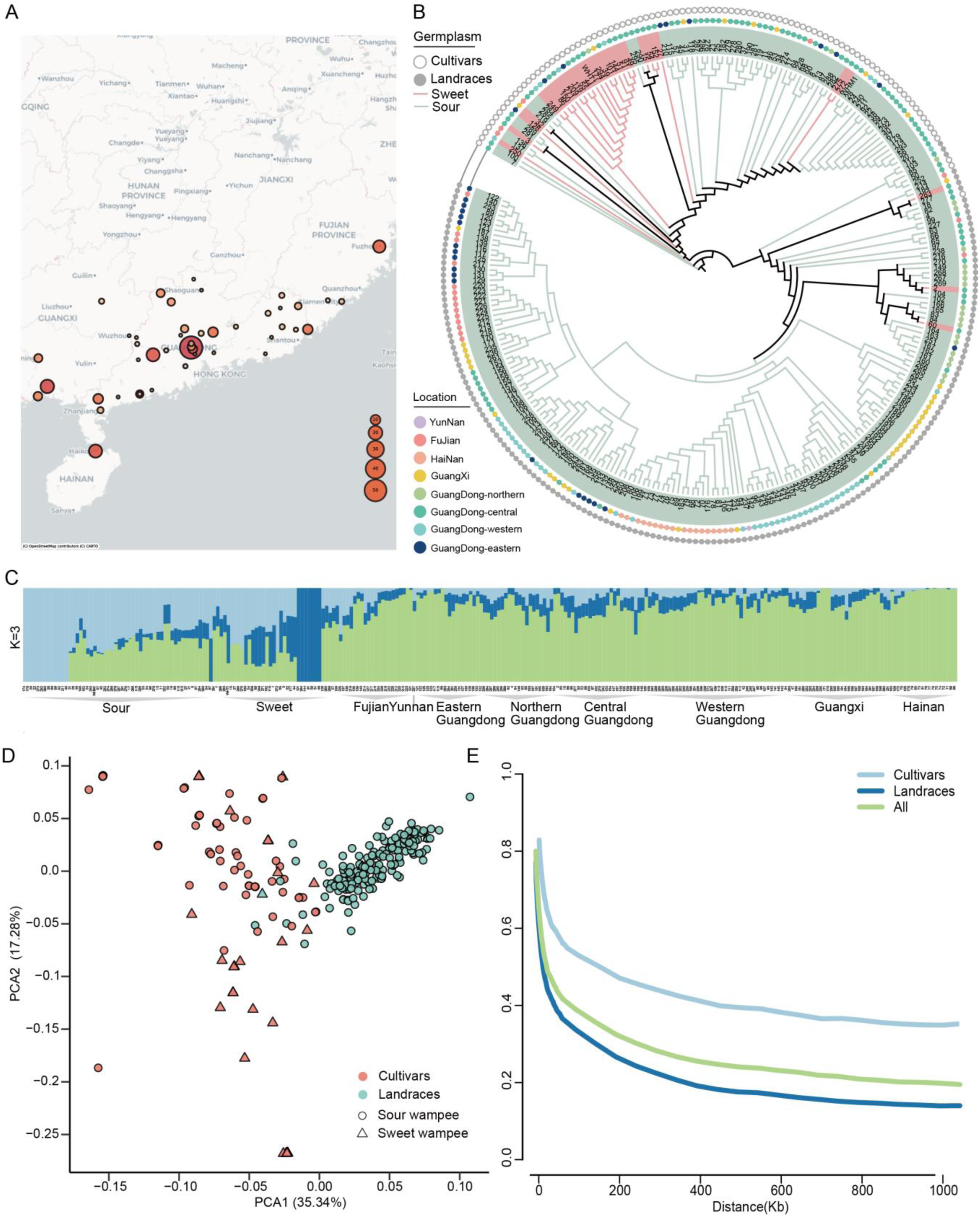
Population structure and genetic diversity in wampees. (A) Geographical distribution of wampee accessions. (B) Maximum likelihood phylogenetic tree of the 266 wampee accessions. (C) Population structure of the 266 wampee accessions inferred using Admixture. The extent of each colored segment indicates the percentage of each genome derived from ancestral populations (K = 3). (D) Principal component analysis (PCA) of 266 wampee accessions. Triangles represent sweet wampees, circles represent sour wampees, light green represents landraces, and light pink represents cultivars. (E) Linkage disequilibrium decay in the 266 wampee accessions, landrace and cultivar accessions.

To further investigate the genetic background of the wampees, we conducted a biogeographical ancestry (admixture) analysis using different ancestral group values (k), with k = 3 yielding the smallest cross-validation error (**Figure S6**), revealing distinct genetic structures within the wampee germplasm from different geographic origins and subpopulations. Specifically, landraces from Hainan displayed nearly a single green ancestry, perhaps because of their geographical isolation from the mainland as well as less social interaction between the island and the main land. In contrast, germplasms from Fujian, Guangdong, and Guangxi showed a more diverse genetic makeup, with multiple ancestries, dominated by the green ancestry. These germplasms had an overall homogenous ancestry, likely due to the geographical proximity of these areas with similar climates and topographies, thus facilitating frequent gene exchanges. Furthermore, certain cultivated sweet and sour wampees exhibited a single ancestry, perhaps because of the repeated selection of individuals with favorable traits through focused selection strategies during the breeding process (**Figure 3C; Figure S7**). PCA showed that the cultivars and landraces were clearly separated, consistent with the phylogenetic tree (**Figure 3D; Figure S8**). Both cultivars and landraces showed rapid linkage disequilibrium (LD) decay (decreasing to 50% of their peak level), with a low baseline (**Figure 3E**). Cultivars showed larger LD pairwise distances than landrace accessions, possibly because of artificial selection leading to reduced genetic diversity in cultivated populations. The π values were comparable in the cultivars (8.55E-04) and landrace accessions (7.33E-04) (**Figure S9 and S10**), whereas their population fixation index (F_ST_) was 0.0433 (**Figure S11**), indicating a relatively minor differentiation between the two subpopulations.

### Selection signals for the sweet and acidic taste of wampee

Fruit flavor is a trait extensively targeted in fruit breeding programs. Wampee varieties can be classified as sweet or sour. To explore the genomic foundation of wampee fruit flavor, we analyzed the genomes of sweet and sour wampees to identify potential selection signatures. The wampee genome was scanned in 100-kb sliding windows moving in 10-kb increments (**Figure 4A**) to locate sections exhibiting elevated allelic divergence and nucleotide diversity in the 266 accessions. For sour wampees, 101 loci were identified, displaying positive selection [Z(F_ST_) ≥1.9; log_2_(π sour/π sweet) ≥ 1.79] (**Figure 4B**), harboring 110 genes (approximately 0.469% of annotated genes) (**Table S13**) with 75 genes containing 182 nonsynonymous SNPs (nsSNPs) (**Table S14**). For sweet wampees, 314 loci were identified in regions displaying positive selection [Z(F_ST_)≥1.9; log_2_(π sour/π sweet) ≤ -1.79] (**Figure 4B**), containing 671 genes (approximately 2.859% of annotated genes) (**Table S13**) with 226 genes carrying 423 nsSNPs (**Table S15**). These nsSNPs likely influence the flavor of wampee fruit by altering protein function. Furthermore, we examined genomic regions displaying high F_ST_ and π ratio (sour/sweet) and identified a 110 kb fragment on Chr1 (43,700,001-43,810,000) containing 11 nsSNPs (**Figure 4C**). The highest F_ST_ and π ratio (sour/sweet) in this region were 6.67 and 2.81 (red points in **Figure 4B**), respectively, indicating a robust selection signal. Within this region, 14 genes were identified, six of which contained nsSNPs (**Figure 4C, Table S16**). Interestingly, the frequency of nsSNPs in the sweet varieties was significantly higher than that in the sour varieties, most of which were homozygous. LD analysis identified sizeable haploblocks within this area, with total SNPs (**Figure S12**) and nsSNPs (**Figure 4C**) demonstrating robust LD values (r^2^), indicating that the homozygous SNPs in this block formed a haplotype likely responsible for the differentiation of wampee fruit flavors.

**Figure 4.**
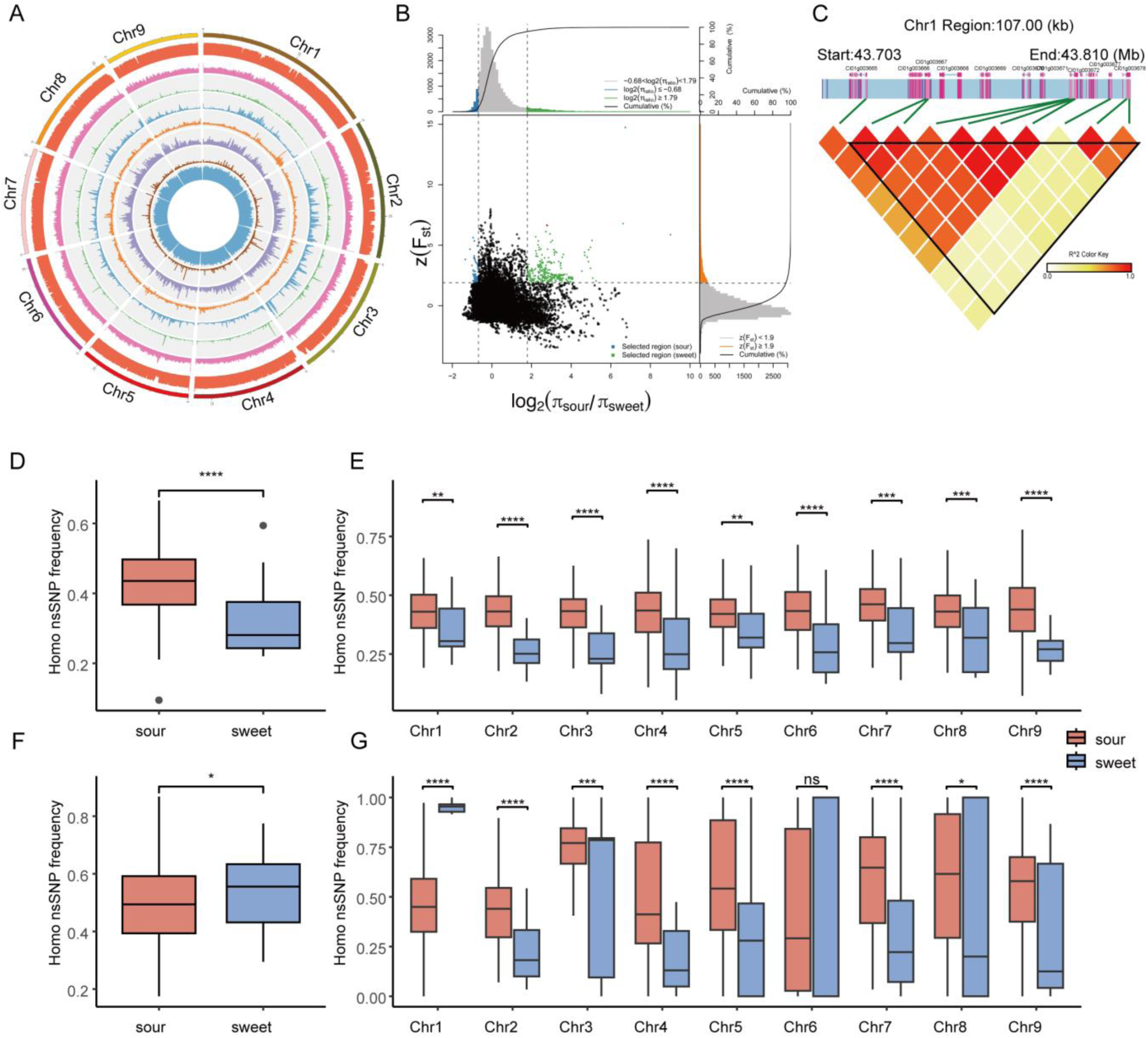
Selection signatures in the wampee genomes considering the differentiation of sweet and sour flavors in fruits. (A) Genetic variation distribution at the genomic level. Outer to inner circles: Density of single-nucleotide polymorphisms (SNPs), insertion-deletions, structural variants, and copy number variants across all chromosomes; fixation index (FST) between sweet and sour accessions, nucleotide diversity, linkage disequilibrium, and neutral evolutionary test (Tajima’s D) within 100-kb frames sliding along the chromosomes. (B) Distribution of FST and π ratios [log_2_(π_sour_/π_sweet_)] for sour and sweet accessions using a 100-kb sliding window and 10-kb steps. Green dots represent genomic regions under positive selection for sour taste; blue dots represent genomic regions under positive selection for sweet taste. (C) Linkage associations of nonsynonymous SNPs within the selective sweep domain on Chr1 (43700001-43810000) aligned with the red markers in panel B. (D-G) Frequency of homozygous nonsynonymous SNPs augmented via selection pressures. (D and F) Box-and-whisker diagrams depicting the incidences across the full genome and the selective locales. (E and G) Box plots contrasting the prevalences across all chromosomes and within the preferential areas of individual chromosomes. Red and blue represent the occurrences in the sour and sweet accessions, respectively. Significant differences were evaluated by conducting a two-tailed *t*-test (ns, not significant; *, **, ***, ****, significance with *P* < 0.05, 0.01, 0.001, and 0.0001, respectively).

The frequencies of homozygous genotype were calculated separately for the sour and sweet accessions. Of the 22,239 identified nsSNPs, 1,789,523 were homozygous, with 1,639,918 and 149,605 in sour and sweet accessions, respectively. The median frequencies of the homozygous genotypes were consistently lower in sweet accessions than in sour accessions at both the genomic and chromosomal levels (**Figure 4D and 4E**). Unexpectedly, the central tendency and upper quartile ranges of hnsSNP frequencies under selection were significantly higher in the sweet accessions (**Figure 4F**). The median hnsSNP frequencies for most chromosomes were lower in sweet accessions than in sour accessions, except for Chr1, where the hnsSNP frequency increased, indicating that the overall increased hnsSNP frequency in the selected regions was mainly driven by the higher frequency in Chr1 (**Figure 4G**). As these nsSNPs showed evidence of adaptive evolution (**Figure 4F**), and the augmented homozygosity suggested altered protein activity resulting from amino acid substitutions. This lends support to the positive role of these nsSNPs in the improvement of sweet wampee fruit flavor.

### Recognition of potential genes participating in sweet and sour traits

Fruit flavor is determined by sugar, acid, and sugar-to-acid ratios ^13^. Organic acids, primarily citric and malic acids, are key to improving fruit quality and flavor ^14^. Therefore, the regulation of fruit acidity is a major goal in wampee research and breeding ^15^. We performed a GWAS based on a multiple loci mixed linear model ^16^ using total fruit acid data collected from 10-15 years-old wampee trees in 2019, 2020, and 2021 (**Table S17**), with a *P* value threshold of 6.03E^-7^ determined utilizing the Genetic Type 1 Error Calculator (v.0.2) ^17^ (**Figure 5A and 5B**; **Figure S13-S14)**. A total of 220 Quantitative Trait Locis (QTLs) were identified, 159 of which were near the gene bodies, and 61 were within the gene sequences (**Table S18**). These QTLs influenced 289 genes (located within 100 kb of the most pivotal SNPs) that were plausible candidates for improving acid accumulation, of which 251 were under selective sweeps and encoded transcription factors, glycosyltransferases, and transport proteins (**Table S18**). The accumulation of organic acids in fruits can be influenced by the sugar content and the interplay between sugar and acid metabolism ^18^. We focused on SNPs with the most significant trait association (Chr1:45211312, -log_10_(P) =12) located in the intergenic region. Thirteen genes were identified within a 50 Kb range upstream and downstream of this SNP. The ethylene response factor (ERF) family gene *ClERF061*, homologous to *ERF061* of sweet orange, had a significantly associated nsSNP (Chr1-45192563) within its coding region, displaying strong LD between Chr1-45192563 and Chr1-45211312 (Figure 5C). Based on the genotypes at Chr1-45192563, the wampee accessions were classified into “C/C” and “G/C” genotypes (Figure 5D). Another gene (Cl01g003921) belonging to the SWEET family was identified as *ClSWEET7*, homologous to *SWEET7* in citrus. Similarly, a nsSNP (Chr1-45517374) significantly associated with *ClSWEET7* was identified showing a strong LD signal (Figure 5G), and the wampee accessions were classified into “T/T” and “C/T” genotype at Chr1-45517374 (Figure 5H). The average fruit organic acid content in the accessions with the homozygous “G/G” or “T/T” genotype exhibited a significantly higher level than those with the heterozygous genotypes (Figure 5D and 5H). Furthermore, the relative expression levels of *ClERF061* and *ClSWEET7* during the color-change period were higher in the sour wampee varieties than in the sweet wampee varieties (*P* < 0.0001) (Figure 5E and 5I). Simultaneously, the association between the relative expression levels of *ClERF061* and *ClSWEET7* and the total acid content/total sugar content was examined. The outcomes displayed a robust positive correlation between the relative expression levels of *ClERF061 or ClSWEET7* and total acid content and a strong negative correlation between the relative expression levels of *ClERF061 or ClSWEET7* and total sugar content (Figure 5F and 5J). We speculated that *ClERF061* or *ClSWEET7* might participate in sugar acid metabolism in wampee fruits; however, proof is still needed from further analyses.

**Figure 5.**
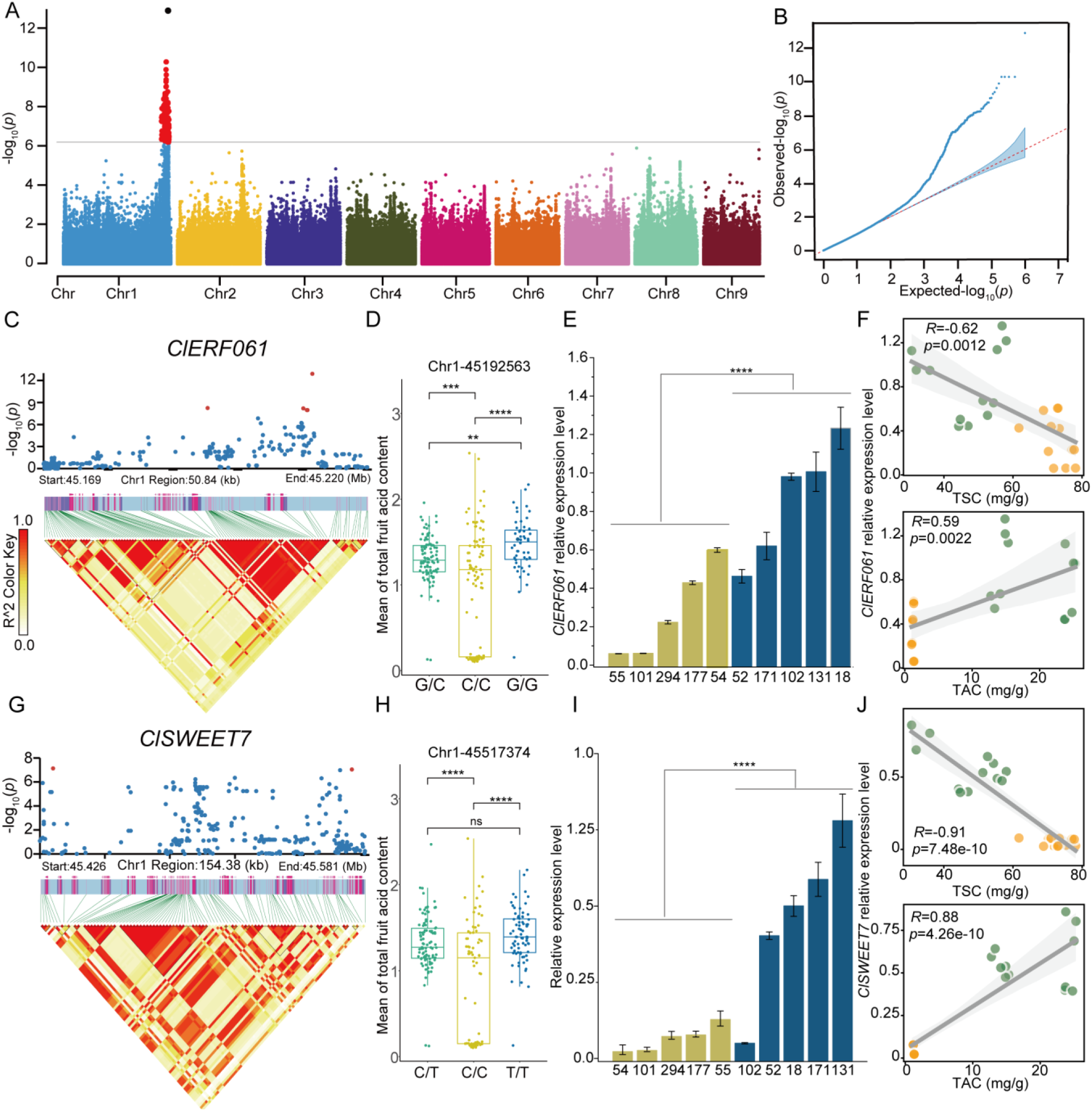
Genome-wide association study (GWAS) recognition of potential target genes involved in sugar-acid metabolism. (A) Manhattan plot based on multiple loci mixed linear model for total acids in fruit. (B) Quantile–quantile plot for the GWAS of total acids in fruit. (C and G) Linkage disequilibrium block diagrams surrounding the statistically notable signals of marker-trait associations. (D, H) Rectangular diagrams of phenotypic information for different genotypes at Chr1-45192563 and Chr1-45517374 loci. The central line of each rectangular diagram symbolizes the median of the data; the upper and lower borders of the box denote the upper and lower quartiles of the data. The circle dots represent the total fruit acid content of each accession. (E, I) Relative gene expression levels of *ClsERF061* and *ClsSWEET7* during the color-change period. Dark yellow and blue columns represent sweet and sour wampee germplasms, respectively. (F, J) Scatterplot of the relative expression of *ClsERF061* and *ClsSWEET7* genes and phenotypes (TSC and TAC); the grey line indicates the linear fit curve, and the shaded area indicates the 95% confidence interval; TSC: total sugar content, TAC: total acid content. Significant differences were determined using a two-tailed *t*-test (ns, not significant; *, **, ***, ****, significance with *P* < 0.05, 0.01, 0.001, and 0.0001, respectively). *Cl01g003921* encodes the bidirectional sugar transporter *ClsSWEET7*. *Cl01g003870* encodes the ethylene-responsive transcription factor *ClsERF061*.

## Discussion

The absence of a high-quality draft genome has been the main hampering determinant in expediting the enhancement of wampee cultivar varieties over the last several decades. In this research, we provide a high-quality chromosome-scale genome assembly of a wampee, which is the most advanced assembly reported to date. Compared to a previous assembly based on 10× genomics technology^11^, our wampee genome assembly showed substantial improvements, with a longer contig N50 of 10.2 Mb and scaffold N50 of 29.2 Mb. This indicated that long read sequencing (LRS) effectively resolved the highly fragmented contigs and scaffolds present in the previous assembly. Additionally, the repeat content in our genome assembly (46%) was higher than that reported by ^11^ (39%), revealing previously undiscovered or low-frequency repetitive sequences, owing to the enhanced resolution provided by LRS. The integration of LRS data significantly enhanced the continuity and completeness of the genome assembly, allowing for better characterization and representation of repetitive regions and complex genomic structures. The high BUSCO score (98%) indicated that the integrity of the present wampee assembly also exhibited improvement compared to that of ^11^ (94.7%). However, extra efforts to attain an exhaustive wampee genome sequence remains desirable.

The evolutionary history of wampee trees remains elusive. Our phylogenomic analysis revealed close affiliations between *C. lansium* and Rutaceae species, including *A. buxifolia*, *P. trifoliata*, *C. sinensis*, and *F. hindsii*. Genome evolution analysis indicated the occurrence of a core eudicot γ triplication event in the wampee genome, which was shared with grape and other core eudicots, but no additional WGD events were found in wampee. Gene family analysis identified 900 expanded and 1,512 contracted gene families, highlighting the impact of evolutionary forces on wampee genetic diversity. Notably, the most closely related KEGG pathway annotations were related to the biosynthesis of alkaloids, considered to be the most important active components with pharmacological effects in wampees. Furthermore, comparative analysis with related and model species is required to identify conserved and differentiated features in these metabolic pathways and to provide insights into their ecological significance in wampees.

The wampee plant originates from southern China and has been grown for thousands of years. We found that the genetic background of wampees was relatively narrow through high-quality genome assembly and population genomics sequencing of wampee varieties, probably because the cultivated species would undergo a domestication bottleneck during artificial selection, resulting in the loss of ancestral SNP diversity. Similar results have been observed for many crops that have experienced domestication bottlenecks, such as wheat and rice. The zygosity of nsSNPs may also carry biological significance in breeding because each nsSNP can induce variances in protein efficacy (Figure 4D), and the homozygosity of activity-altering nsSNPs will further enhance or diminish these activity changes. In our study, the frequency of sweet-type homozygous nsSNPs was significantly increased in the selected regions, implying selection during the domestication of wampees and fixation of nsSNPs in a homozygous state. Sour wampee is the predominant known wampee variety, whereas sweet wampee is a minority, suggesting that wampee undergoes a process of shifting from sour to sweet during artificial breeding, similar to the breeding process of other fruit trees.

A GWAS analysis utilizing the total acid content data of wampee fruit obtained over three consecutive years was conducted, identifying 220 marker-trait associations, with 10 nsSNPs situated within the coding domains of relevant genes. Notably, the traits were associated with 289 genes. Sugar and acid metabolism in fruit are closely interconnected and mutually influence and regulate each other during fruit development and ripening processes ^18^, where certain sugar molecules can serve as precursors for acid synthesis or act as regulatory factors in the acid synthesis process. GWAS identified a significant QTL locus for organic acid content containing the gene *Cl01g003921*, which encodes SWEET7. SWEET, a bidirectional sugar transporter ^19^, may be involved in the transportation of sugars during fruit development and ripening stages in multiple crops, such as tomato, grape, apple, sugar apple, and citrus. In tomatoes, the *SISWEET7a* gene exhibits the highest relative expression levels in the pedicel and fruit vascular bundles during the ripening period, indicating its regulatory role in fruit development^20^. Similar results are observed in sugar apples (*Annona squamosa*), where the *SWEET16-1* gene shows the highest expression in the pedicels of mature fruits, suggesting its role in fruit development ^21^. Additionally, the grape SWEET10 protein mediates sugar accumulation in fruits ^22^. In apples, the sugar transport protein MdSWEET9b contributes to sugar accumulation, where MdSWEET9b specifically transports sucrose, facilitating regular yeast development and total sugar content ^23^. In *Hylocereus undatus*, HuSWEET12a/13d regulates fruit sugar accumulation, and transgenic plants overexpressing these genes show higher levels of sugar accumulation ^24^. The activation of CitSWEET6 expression in Wenzhou honey orange promotes fructose accumulation ^25^. In citrus, overexpression of CitSWEET11d significantly increased sucrose content in callus tissue, with CitERF16 exhibiting a similar expression pattern to CitSWEET11d during fruit development ^26^. In addition, CitERF16 was positively aligned with sucrose content and promoted sucrose accumulation by regulating CitSWEET11d.

Another candidate gene identified via GWAS, *Cl01g003870*, encodes an ethylene-responsive transcription factor belonging to the APETALA2 (AP2)/ERF superfamily of plants. ERFs demonstrate a wide scope of biological functions, participating in both stress responses and growth and developmental processes, including controlling color, texture, flavor, and other important fruit quality traits. The main function of ERFs is achieved through direct regulation of target genes, although some may act by regulating other transcription factors. A typical ERF domain protein, VvDREB, has been identified in grapes ^27^. It interacts with VvMSA to regulate the glucose transporters, thereby affecting the sweetness of grapes. DkERF10 and DkERF22 have been identified in *Diospyros kaki*, which can regulate the promoters of *DkADH1* and *DkPDC2*, which participate in the tannin metabolism of persimmon fruit, thus influencing astringency removal in persimmons ^28^. Thus, our GWAS candidate genes included genes from the AP2/ERF and SWEET protein families.

In conclusion, we report a chromosome-level genome assembly for the wampee cultivar ’JinFeng’ and investigate the population genetic diversity of wampees via population sequencing analysis of 266 wampee accessions, demonstrating population hybridization and gene flow among cultivars from major wampee growing areas. These findings provide a research direction for the future improvement of wampee flavor by offering numerous promising candidate genes for enhancing the desired flavor traits of wampee by integrating genomic, transcriptomic, and metabolomic data.

## Materials and methods

### Plant materials

The *Clausena lansium* (Lour.) Skeels. cv. ‘JinFeng’ originated in Guangzhou of Southern China, which exhibited relatively uniform ripening characteristics and excellent fruit quality, were well-suited for cultivation not only in Guangdong Province but also in other major wampee-producing regions across China. An 8-year-old *Clausena lansium* (Lour.) Skeels tree named Jinfeng from the Wampee Germplasm Repository affiliated to Ministry of Agriculture and Rural Affairs was used for genome sequencing and *de novo* assembly. Total genomic DNA was isolated from fresh young leaves utilizing the CTAB (Cetyltrimethylammonium Bromide) reference method. The concentration of DNA was measured with the Qubit® DNA Assay Kit in a Qubit® 2.0 Fluorometer (Life Technologies, USA). For RNA extraction, samples were collected from six different tissues, including root, leaf, stem, flower, fruit, and seed from the same wampee tree. Total RNA was obtained applying the OMEGA RNA extraction method (OMEGA, USA).

### Library construction and sequencing

Genomic DNA was used in library preparations utilizing the NEB Next® Ultra DNA Library Prep Kit for Illumina® (NEB, USA) following the recommended protocols. Initially, DNA was randomly degraded, and then the 3’ ends of DNA fragments were adenylated and end-repaired using the NEB Next Adaptor with a hairpin loop structure. Paired-end DNA fragments of 125/150/250 bp were produced and sequenced on an Illumina HiSeq platform. For the single molecule real-time (SMRT) library preparation, the SMRTbell Template Prep Kits were utilized according to the instructions. Briefly, DNA fragments with a target size of approximately 20 kb (selected from > 15 kb size) underwent successive treatments, including damage repair, end repair, blunt-end adaptor ligation, and size selection. Finally, the 20kb SMRTbell libraries were sequenced using the PacBio Sequel platforms at Novogene company (Beijing, China). Hi-C raw reads obtained from the Illumina platform were utilized to construct chromosome-level scaffolds, with appropriate filtering applied before analyses. These reads were mapped to the assembled contigs using BWA (v0.7.17) with default parameters ^29^ which enabled the establishment of contact information among the contigs. Ultra-long-range scaffolding of the de novo genome assemblies were performed using LACHESIS (v201701) software ^10^, leveraging the genomic proximity signals provided by the Hi-C data. To assess the quality of the assembled genome, BUSCO was employed ^30^.

### Genome assembly and quality assessment

Genome size and heterozygosity estimation for *C. lansium* were conducted using Jellyfish (v2.1.3) with a k-mer size of 17 on Illumina short reads ^31^. Genome assembly and obtain chromosome-level scaffolding was conducted using a combined approach utilizing Illumina, PacBio, and Hi-C data. From the raw PacBio reads, sequences with adaptor sequences or of poor quality were excluded. The remaining reads underwent self-correction using Falcon (v0.7) ^8^, and consensus sequences were generated from these error-corrected reads. To improve assembly accuracy, the consensus sequences were further polished with Illumina short reads using nextpolish (v1.1.0) ^9^.

### Repeats and gene annotation

A comprehensive annotation for *C. lansium* genome was conducted to identify repetitive elements and structural features. For repeat annotation, a combination of homology-based alignment and *de novo* prediction methods were utilized. For Homology alignment, interspersed repeats were recognized by matching against the Repbase database ^32^ using RepeatMasker and RepeatProteinMask tools (http://www.repeatmasker.org/). For *de novo* prediction, a repeat library was built with LTR_Finder (http://tlife.fudan.edu.cn/ltr_finder/) ^33^, RepeatScout (http://www.repeatmasker.org/), and RepeatModeler (http://www.repeatmasker.org/RepeatModeler.html) with default parameters. Subsequently, the identified repeats were masked using RepeatMasker (http://repeatmasker.org/). The structural annotation of the genome was established through a dual approach of homology-based prediction and RNA-Seq assistance. Homologous protein sequences from Ensembl, NCBI, and other sources were aligned to the genome using TblastN (v2.2.26; E-value ≤ 1e^−5^) ^34^. Accurate spliced alignments were generated using GeneWise software (v2.4.1) ^35^ by comparing the matching proteins to the homologous genome sequences. Additionally, an automated gene prediction pipeline incorporated *ab initio* gene prediction methods such as Augustus (v3.2.3) ^36^, Geneid (v1.4) ^37^, Genescan (v1.0), GlimmerHMM (v3.04) ^38^, and SNAP^39^.

To further aid genome annotation, transcriptome reads were assembled with Trinity (v2.1.1) ^40^. RNA-seq reads from various tissues were aligned to the genome fasta using Hisat (v2.0.4) with default parameters to identify exon regions and splice positions ^41^. The alignment results were used for genome-based transcript assembly using Stringtie (v1.3.3) with default parameters ^42^. A non-redundant reference gene set was generated by merging genes predicted through multiple methods, including EvidenceModeler (EVM, v1.1.1) with PASA (v2.1) ^43^, terminal exon support, and incorporating masked transposable elements as input. Gene roles were designated by matching protein sequences to the Swiss-Prot database^44^ using Blastp (with a threshold of E-value ≤ 1e^−5^). Annotation of motifs and domains was performed using InterProScan (v5.31) ^45^ and publicly available databases such as ProDom (http://prodom.prabi.fr/prodom/current/html/home.php), PRINTS (http://130.88.97.239/PRINTS/index.php/), Pfam (https://pfam.xfam.org/), SMRT (http://smart.embl-heidelberg.de/), PANTHER (http://pantherdb.org/), and PROSITE (http://www.expasy.org/prosite/). Gene Ontology (GO) IDs were assigned based on the corresponding InterPro entry. Protein functions were predicted by transferring annotations from the closest BLAST hit (E-value <10^-5^) in the SwissProt and NR databases. Additionally, gene sets were mapped to the Kyoto Encyclopedia of Genes and Genomes (KEGG, http://www.genome.jp/kegg/) pathway to determine optimal match for each gene^46^. Non-coding RNA annotations, including tRNA, rRNA, miRNA, and snRNA, were conducted. tRNAs were estimated utilizing the tRNAscan-SE program (http://lowelab.ucsc.edu/tRNAscan-SE/) ^47^ with relative species’ rRNA sequences as references. Other ncRNAs, such as miRNAs and snRNAs, were discovered through a search against the Rfam database (http://rfam.xfam.org/) ^48^ using infernal software with default parameters (http://infernal.janelia.org/) ^49^.

### Gene family and phylogenetic analysis

A comparative analysis of protein sequences from *C. lansium* and 16 related species (*C. sinensis*, *P. trifoliata*, *F. hindsii*, *A. buxifolia*, *D. longan*, *E. japonica*, *Theobroma cacao*, *Arabidopsis thaliana*, *V. vinifera*, *Populus trichocarpa*, *Oryza sativa*, *Averrhoa carambola*, *Pistacia vera*, *Aquilegia coerulea*, and *Eucalyptus grandis*) was constructed to investigate duplication events and classify species-specific genes. Genome sequences of the related species were sourced from the National Center for Biotechnology Information (NCBI) Database, with the exception of *P. trifoliata*, *F. hindsii*, and *A. buxifolia*, which were obtained from the Citrus Pan-genome to Breeding Database (CPBD).

Gene families were clustered using OrthoMCL (RRID:SCR 007839) ^50^ and single-copy gene family prediction was performed. An alignment was conducted for each gene family utilizing Muscle (http://www.drive5.com/muscle/) ^51^, and positions with ambiguous alignments were removed with Gblocks (http://molevol.cmima.csic.es/castresana/Gblocks.html) ^52^. The Neighbor-joining tree was constructed using RAxML v8.2.12. (https://github.com/stamatak/standard-RAxML) ^53^. To estimate divergence times, the MCMCtree program (http://abacus.gene.ucl.ac.uk/software/paml.html) integrated into PAML ^54^ was utilized. To analyze gene family evolution as a stochastic birth and death process, a likelihood model from the Cafe software package (http://sourceforge.net/projects/cafehahnlab/) was employed ^55^. This model considers gene family expansion or contraction per gene per million years, independently along each branch of the phylogenetic tree. The significance of changes in gene family size on each branch was inferred, considering the phylogenetic tree topology and branch lengths.

### Whole-genome resequencing and variation calling

A collection of 266 wampee individuals were chosen for resequencing (see Data Table S17). The raw reads were processed to remove reads with adapters (Phred score < 5 or N > 10%). The clean reads obtained after quality control and adapter removal were then mapped to the compiled reference genome using BWA (v0.7.17) ^29^, and the mapping statistics were assessed. PCR duplicates were marked and sequences were realigned using Picard v3.0.0 (https://github.com/broadinstitute/picard). Coverage and sequencing depth were calculated based on the final BAM files using SAMtools v1.9 ^56^. The Genome Variant Call Formats (gVCFs) were obtained for each individual using freebayes ^57^. These gVCF files were merged to generate the VCF file for all individuals using Bcftools v1.17 ^56^. The filtering parameters of --minDP 2, --maxDP 1000, --minQ 30, --min-alleles 2, and --max-alleles 2 with VCFtools v0.1.16 ^58^ were applied to obtain a total of 1,040,809 SNPs. For the population study, the VCF file was refined through filtering to construct the basic set of SNPs by excluding non-biallelic SNPs and those with a max missing rate > 0.9. Additionally, SNPs with minor allele frequency (MAF) < 0.01 were removed. The obtained SNPs were then annotated with SnpEff v5.1 ^59^ for further analysis.

### Population genetic and Phylogenetic analysis

The population structure analysis was performed using the maximum-likelihood approach implemented in ADMIXTURE v1.3.0 ^60^. Individual-based clustering analysis was carried out, and cross-validation was employed to convergence. The minimum cross-validation error was utilized to determine the optimal number of clusters (K: 2–10). High-quality SNPs were selected to construct the neighbor-joining tree using PLINK v1.90b6.21 ^61^ and IQ-TREE ^62^. The neighbor-joining tree was visualized using the interactive tree of life (iTOL, https://itol.embl.de) tool. The PCA analysis for all SNPs was carried out using PLINK v1.90b6.21 and visualized through ggplot2 package in the R environment.

### Linkage disequilibrium analysis

The mean squared correlation coefficient (r^2^) values between any two SNPs within a 500 kb was calculated using PopLDdecay v3.41 software ^63^ to estimate and evaluate the patterns of LD decay in each population.

### Selective sweep analysis

*FST*, nucleotide diversity (θπ), and Tajima’s D for different populations were computed in a sliding window of 100 Kb with a step size of 10 Kb using VCFtools v0.1.16 ^58^. *Fst* was employed to assess the level of genomic divergence between two subgroups. θπ was utilized to estimate the genomic diversity within each subgroup. Next, the θπ ratios were analyzed for each sliding window for the two subgroups. By designating the top 5% of the log-odds ratios for both θπ and *Fst*, putative selection targets were identified. The sour and sweet accessions were analyzed to identify loci that experienced selective sweeps.

### Genome-wide association study

The total acid content of wampee fruit was obtained from 2019 to 2021.Titratable acid content was quantified based on the previous publication ^64^ with an Automatic Potentiometric Titrator (TITRALAB TIM840, Loveland, Colorado, USA) and was expressed as “g/100 g FW”. GWAS analysis with different models (MLMM, Blink, ECMLM, FarmCPU) of GAPIT3 ^65^ was conducted using high-quality SNPs and pretreated traits of total acid content in fruits pretreating with the Best Linear Unbiased Prediction (BLUP) model. The P-value threshold for significance (6.03×10−7) was determined based on a uniform threshold of 1/n, where n represents the effective number of independent SNPs calculated using the Genetic type 1 Error Calculator (v.0.2) ^17^. The results were presented graphically using the CMplot package in the R environment. Additionally, the regional haplotype blocks were displayed using the LDBlockshow tool ^66^.

## Acknowledgments

We thank Chunyu Li (Institute of Fruit Tree Research, Guangdong Academy of Agricultural Sciences) for useful advice on the experiment design. We thank Dr. Ming Yan, Dr. Dilay Ayhan Hazal, Jie Sun, Tan Meng for their assistance in data analysis. This work was supported by the Special Fund for Agricultural Industry Revitalization Project under Rural Revitalization Strategy (2022-NPY-00-036), the Special Funding Project for Scientific and Technological Talents Introduction at Guangdong Academy of Agricultural Sciences (R2021YJ-YB3022), the Innovation Team of Modern Agricultural Industry Technology System in Guangdong Province of China (2022KJ116, 2023KJ116), the National Tropical Plants Germplasm Resources Center (2023), the Guangzhou Basic and Applied Basic Research Foundation (2023A04J0141), and Shandong Provincial Science and Technology Innovation Fund. L.G. is also supported by Taishan Scholar Program and Natural Science Foundation for Distinguished Young Scholars (ZR2023JQ010) of Shandong Province.

## Contributions

H.Q.C., L.G. and Y.S.L. designed the study; Z.C., J.S.Q., C.P., X.X.C. and Y.S.L. collected and processed the wampee samples; H.Q.C., Z.C., J.X.W., X.F.W. and C.P conceived experiment and analyzed the data; H.Q.C., L.G. and J.X.W performed the result interpretation; H.Q.C., J.X.W., B.W.Y., X.R.W. prepared figures and tables; H.Q.C. and L.G. wrote the paper. All authors read and approve the manuscript.

## Data availability statement

The genome sequencing data, RNA-seq data, Hi-C data, and genome assembly of *C. lansium* produced in this research have been submitted to the Genome Sequence Archive (GSA) database at the National Genomics Data Center of China National Center for Bioinformation, with the accession number PRJCA022284.

## Supplementary information

**Figure S1.** Chromosome DAPI staining of wampee samples (bar = 5 μm).

**Figure S2.** K-mer analysis of wampee genome characteristics.

**Figure S3.** Genome comparison between *Clausena lansium* (Chr1∼9) and *Citrus sinensis* (Chr1∼9), *Vitis vinifera* (Chr1-18), and *Eriobotrya japonica* (Chr1-8), with whole-genome duplication events highlighted in red boxes.

**Figure S4.** SNP density of chromosomes.

**Figure S5.** Distribution of SNP mutation types in the wampee accessions.

**Figure S6.** Annotation of SNP mutations.

**Figure S7.** CV plot in wampee accessions using Admixture (K = 2-10).

**Figure S8.** Population structure of 266 wampee accessions inferred using Admixture. Each colored segment’s length represents the proportion of each genome inferred from ancestral populations (K = 2-10).

**Figure S9.** The percentage variance-explained values of each component in the PCA results.

**Figure S10.** Genome-wide nucleotide diversity (π) of cultivar accessions on the nine chromosomes.

**Figure S11.** Genome-wide nucleotide diversity (π) of landrace accessions on the nine chromosomes by ggplot2 R Packages.

**Figure S12.** Genome-wide F_ST_ between landrace and cultivar accessions on the nine chromosomes. Dotted line, 1% threshold.

**Figure S13.** Linkage disequilibrium of all SNPs in the selective region on Chr1 (43700001-43810000) corresponding to the red dots in Figure 4B.

**Figure S14.** Quantile–quantile plot displaying the GWAS results for total acid content in wampee fruit using: A, Blink; B, Compressed Mixed linear Model (CMLM); C, Enriched Compressed Mixed Linear Model (ECMLM); D, Fixed and random model Circulating Probability Unification (FarmCPU) models.

**Figure S15.** Manhattan plot displaying the GWAS results for total acid content in wampee using FarmCPU, MLMLM, Blink, CMLM, and ECMLM models. The threshold for significance is *P* < 6.03E^-7^ (-log_10_*P* > 6.22).

**Supplemental Tables 1-18.**

