## Supplementaey Figure for "Wampee chromosome-level reference genome elucidates fruit sugar-acid metabolism"

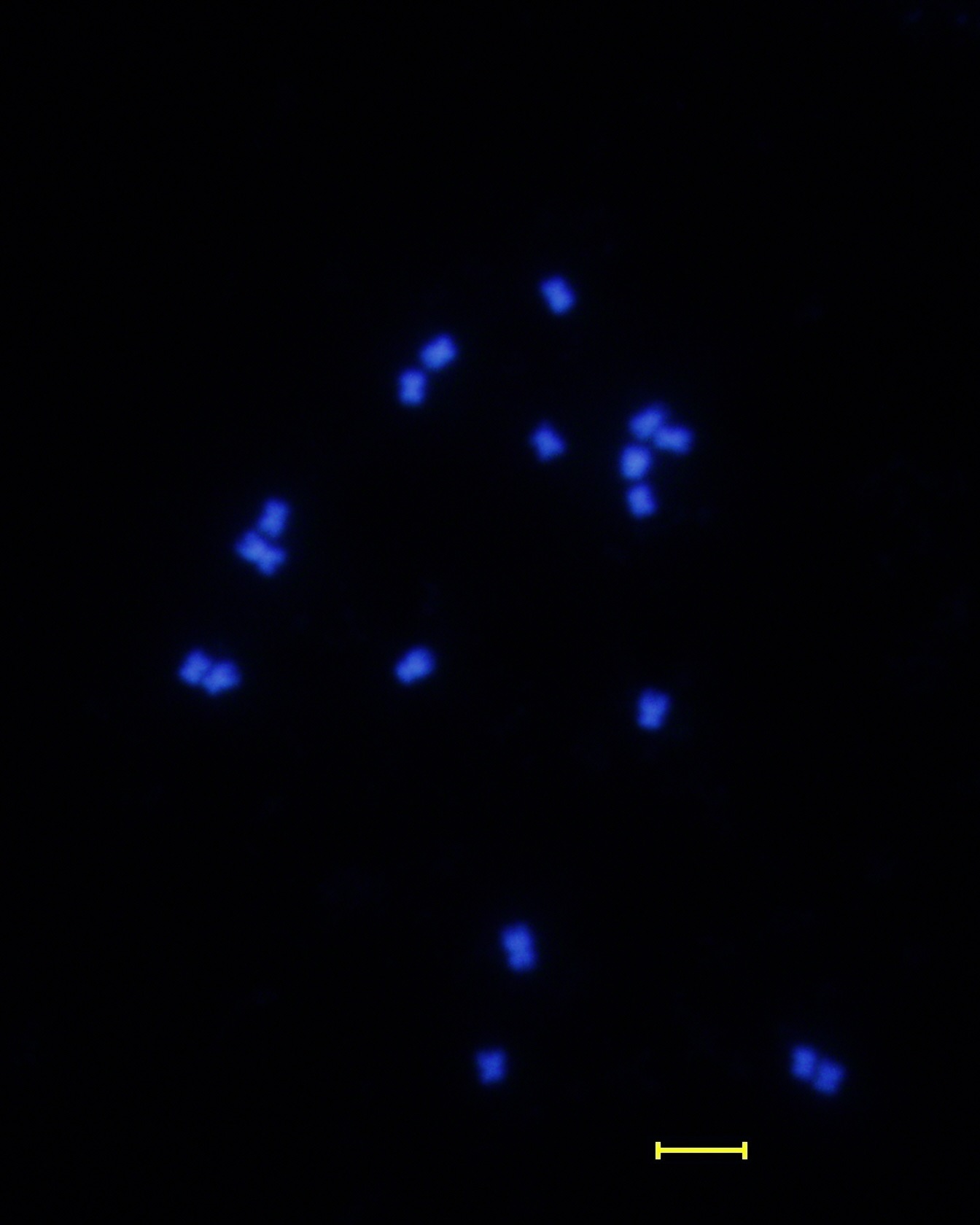

Figure S1. Chromosome DAPI staining of the wampee sample. (Bar: 5 μm)

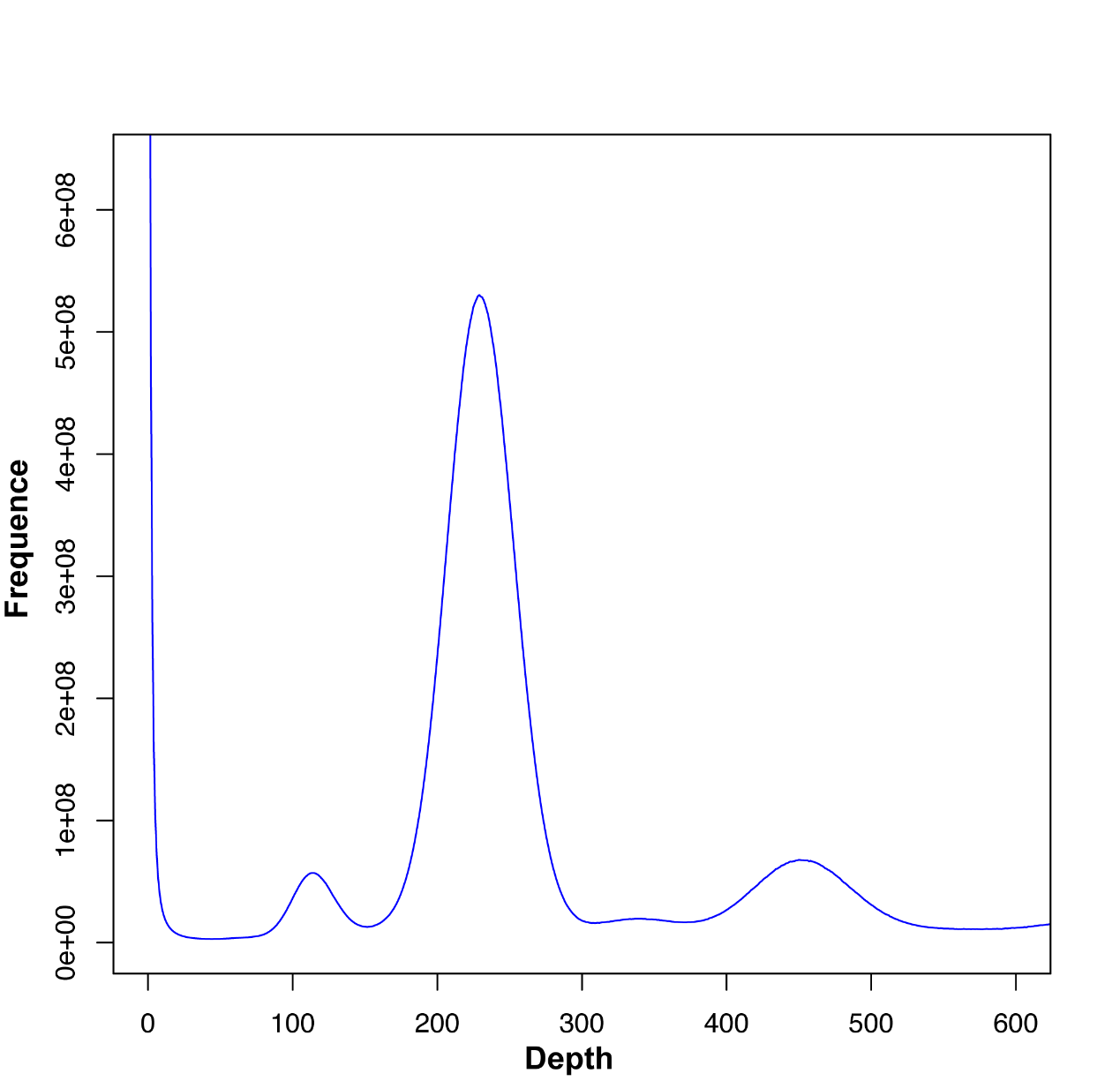

Figure S2. The k-mer analysis of wampee genome characteristics.

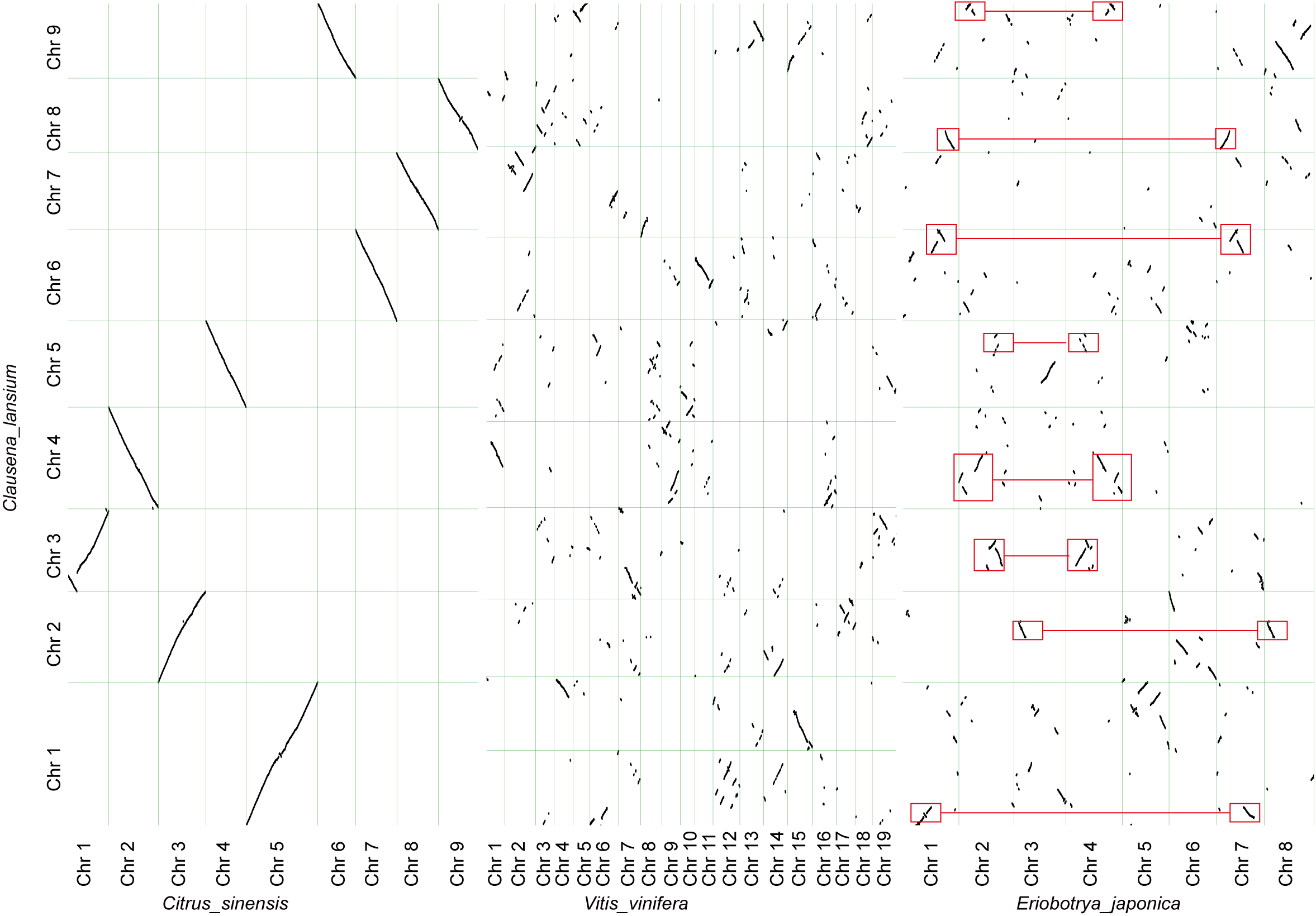

Figure S3. Genome comparison between *C. lansium* (Chr1~9) and *Citrus sinensis* (Chr1~9), *Vitis vinifera* (Chr1-18), *Eriobotrya japonica* (Chr1-8) respectively, with the whole genome duplication events highlighted with red box.

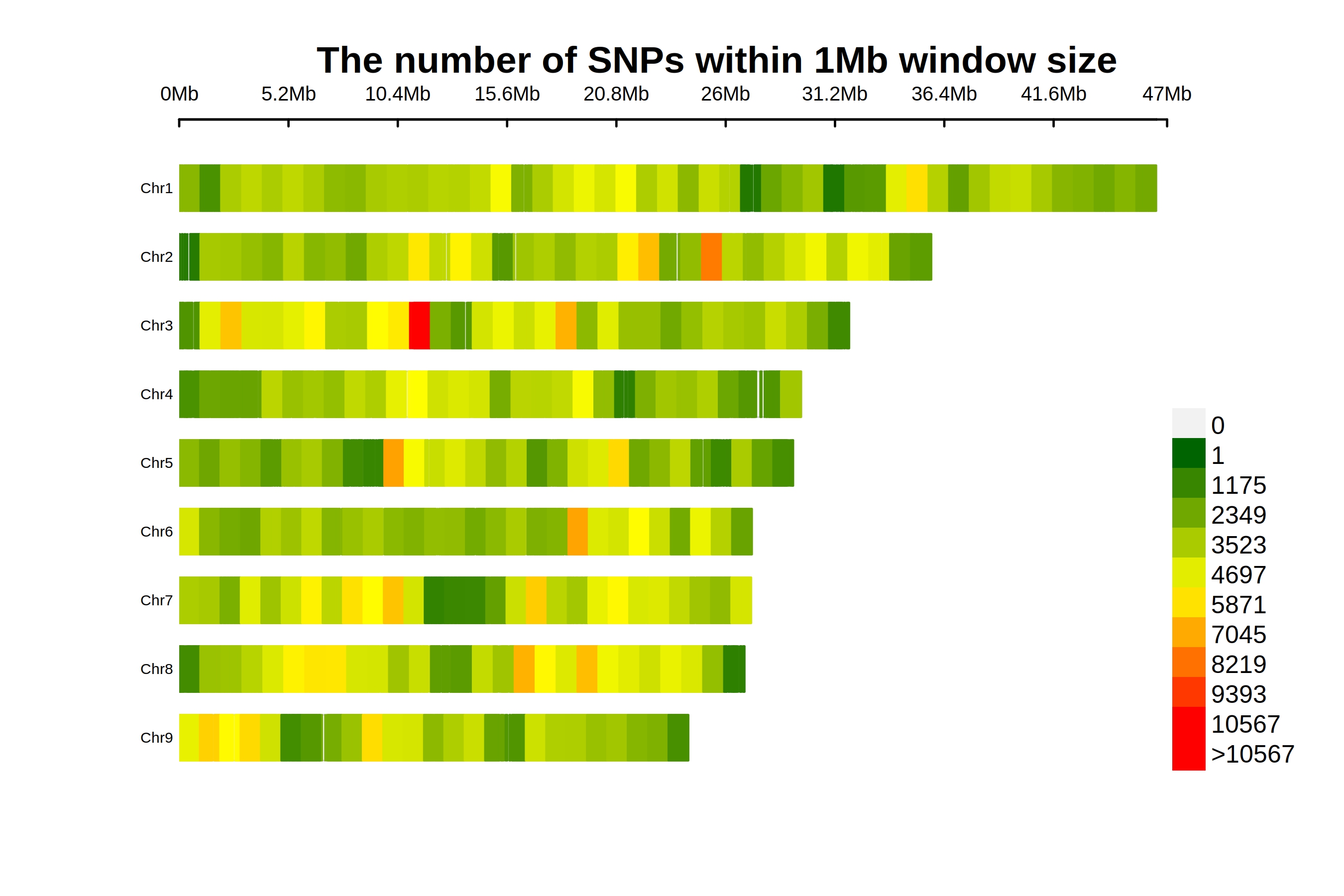

Figure S3. SNP density of chromosomes.

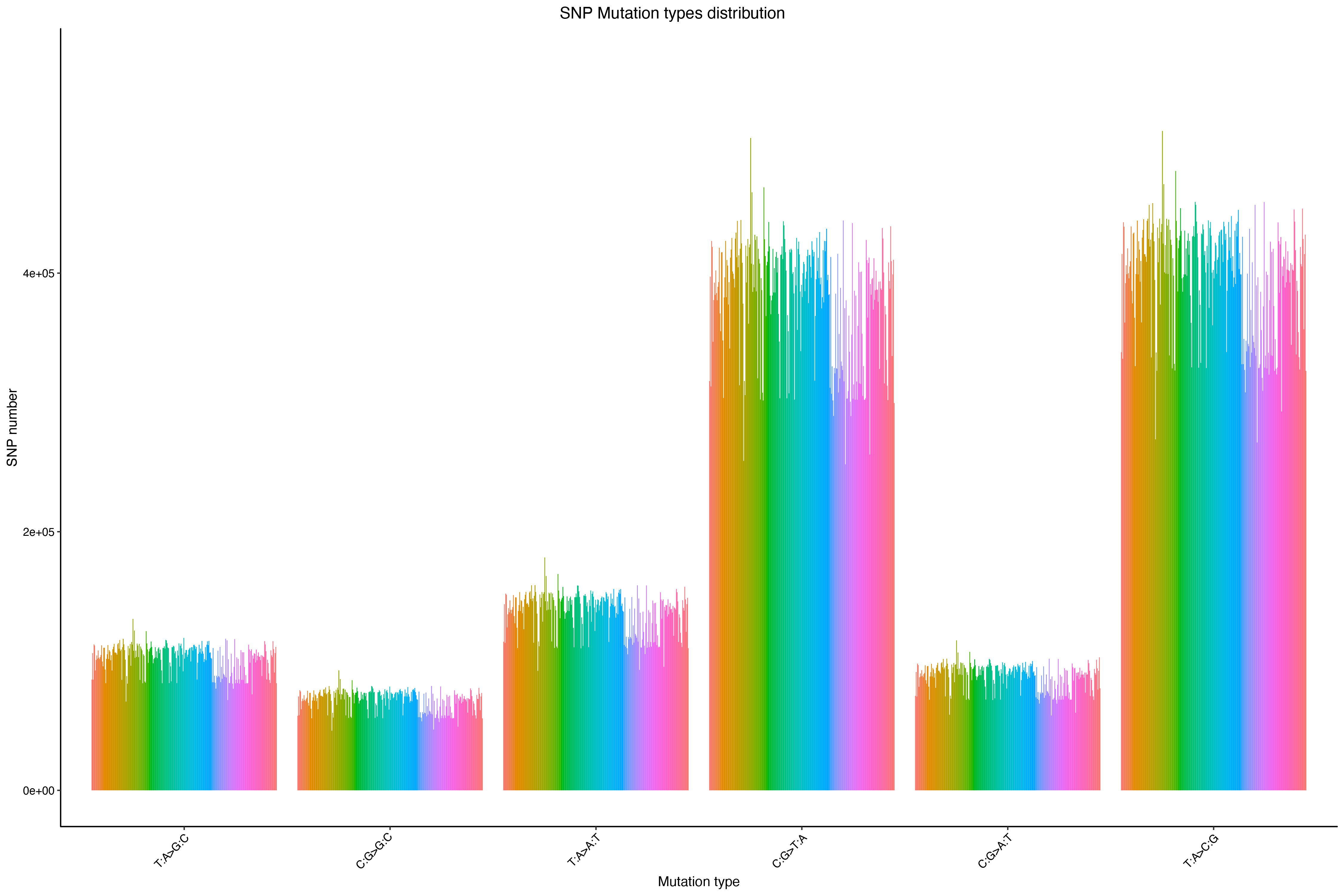

Figure S4. The distribution of SNP mutation types in the wampee accessions.

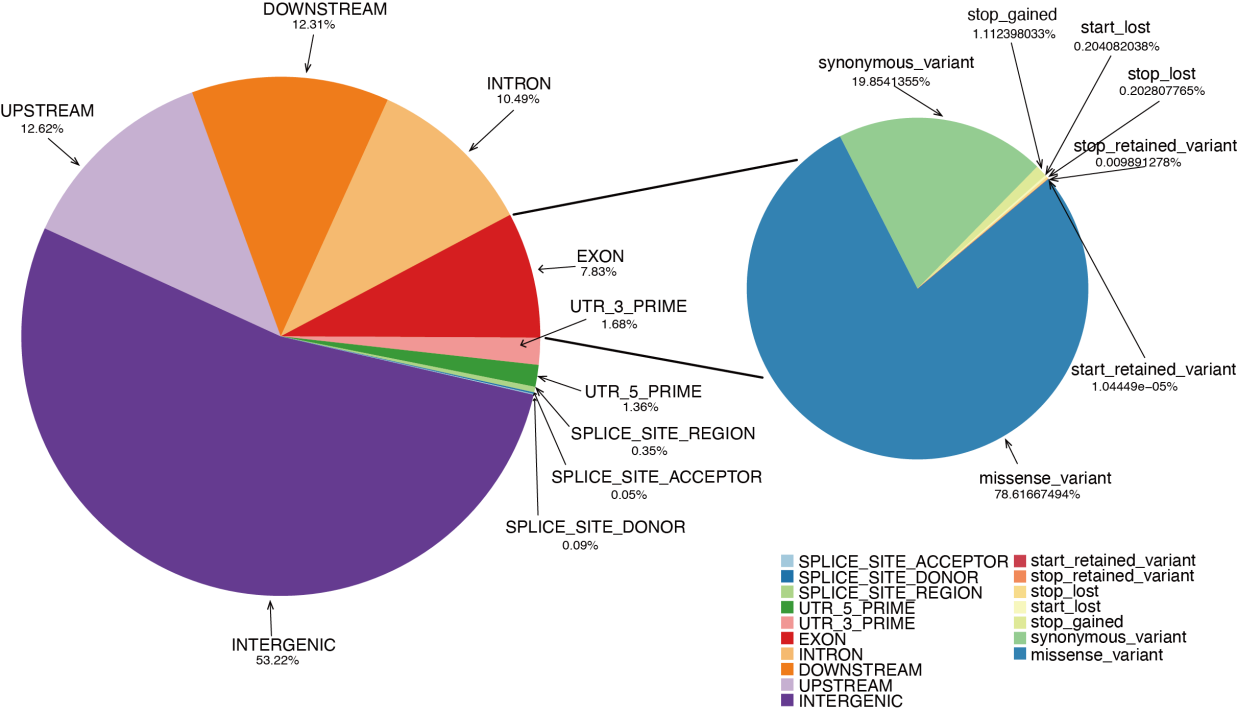

Figure S5. The annotation of SNP mutations.

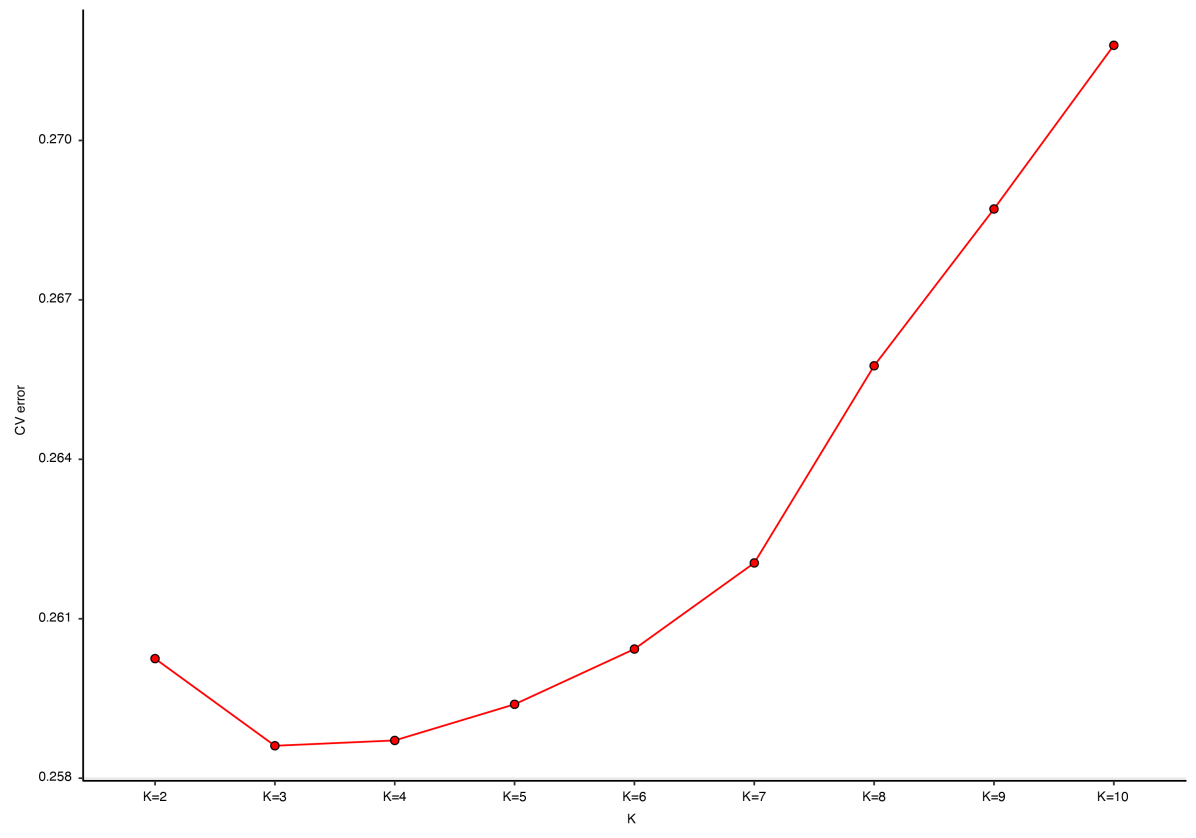

Figure S6. The (CV) plot in wampee accessions using Admixture with K value ranging from 2 to 10.

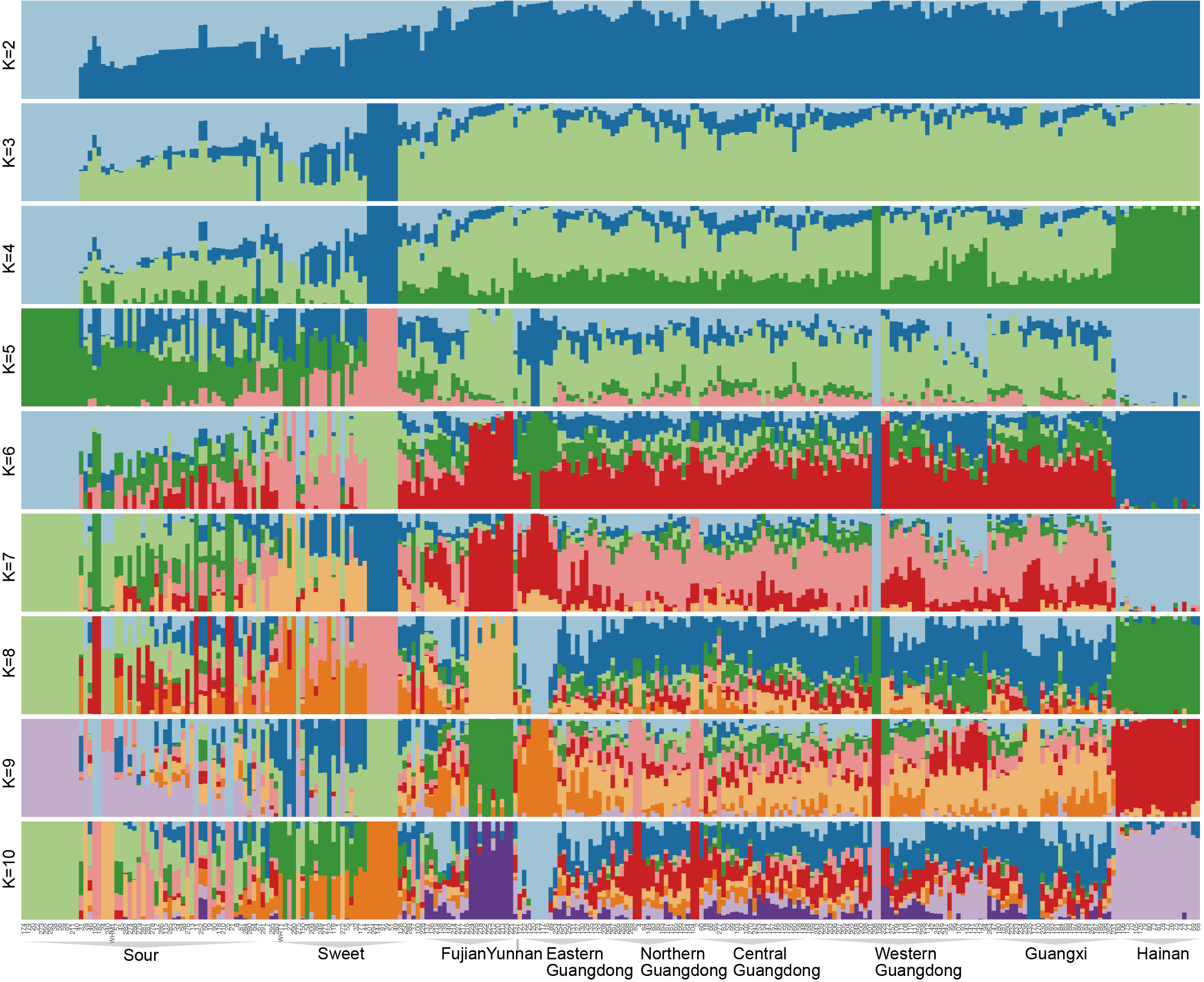

Figure S7. Population structure of 266 wampee accessions inferred by Admixture. The length of each colored segment represents the proportion of each genome inferred from ancestral populations (K = 2 - 10).

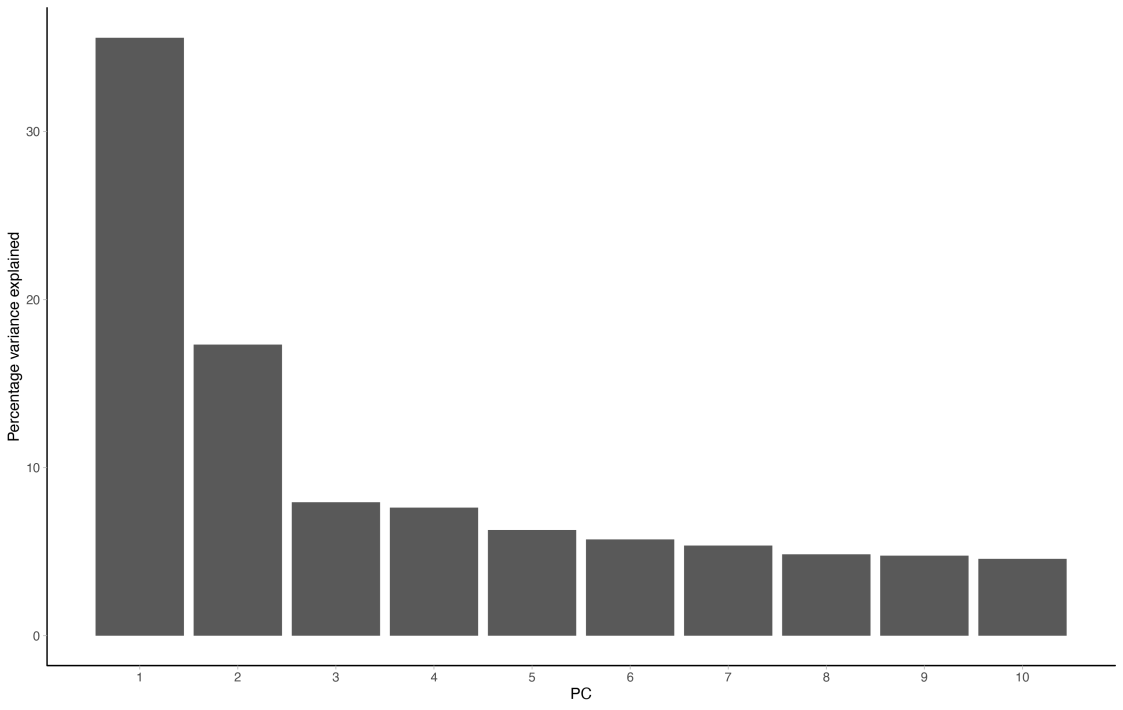

Figure S8. The percentage variance explained values of each component from PCA results.

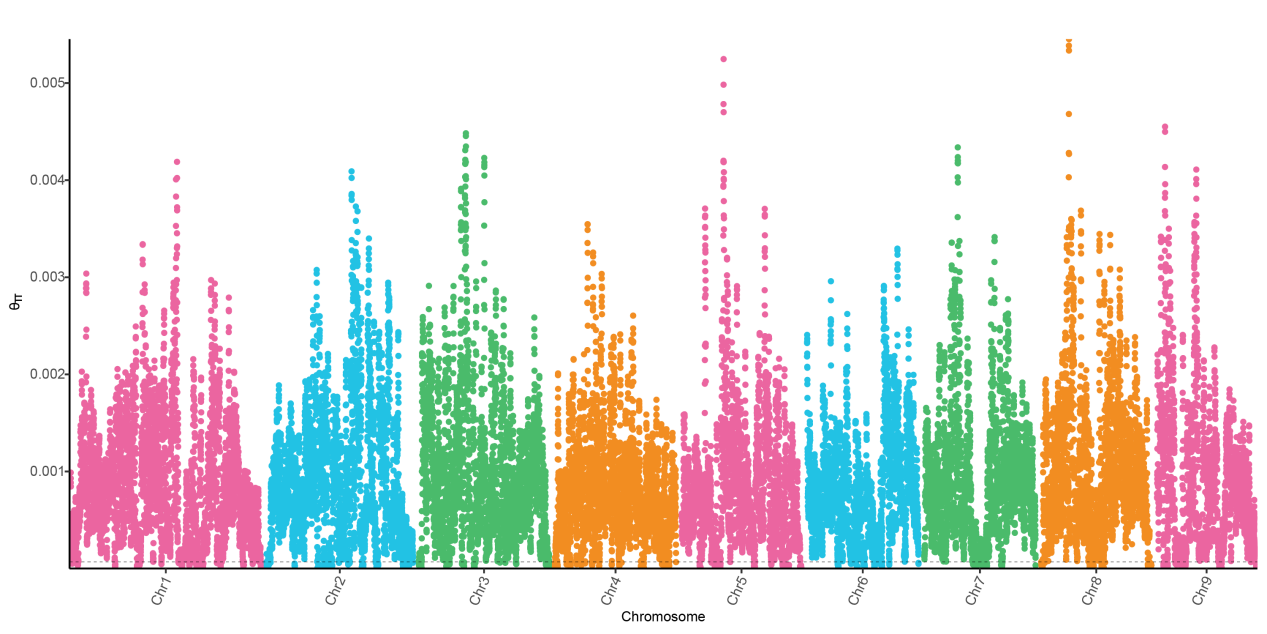

Figure S9. Manhattan plot of genome-wide nucleotide diversity (π) of cultivar accessions on the 9 chromosomes.

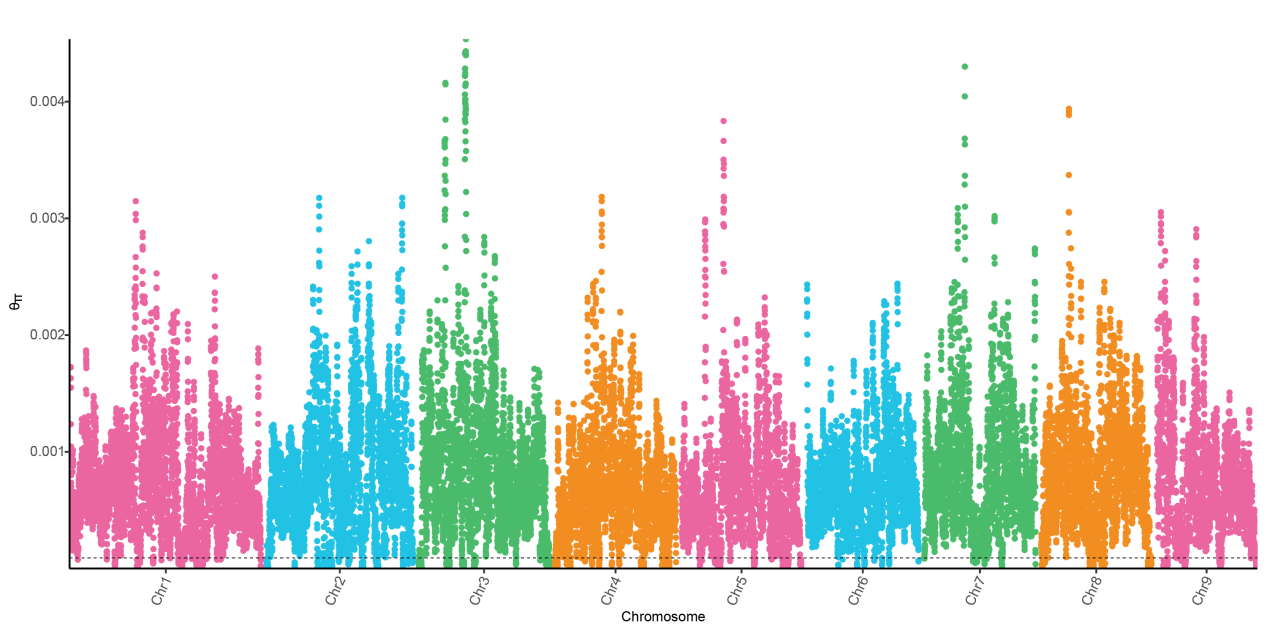

Figure S10. Manhattan plot of genome-wide nucleotide diversity (π) of landrance accessions on the 9 chromosomes.

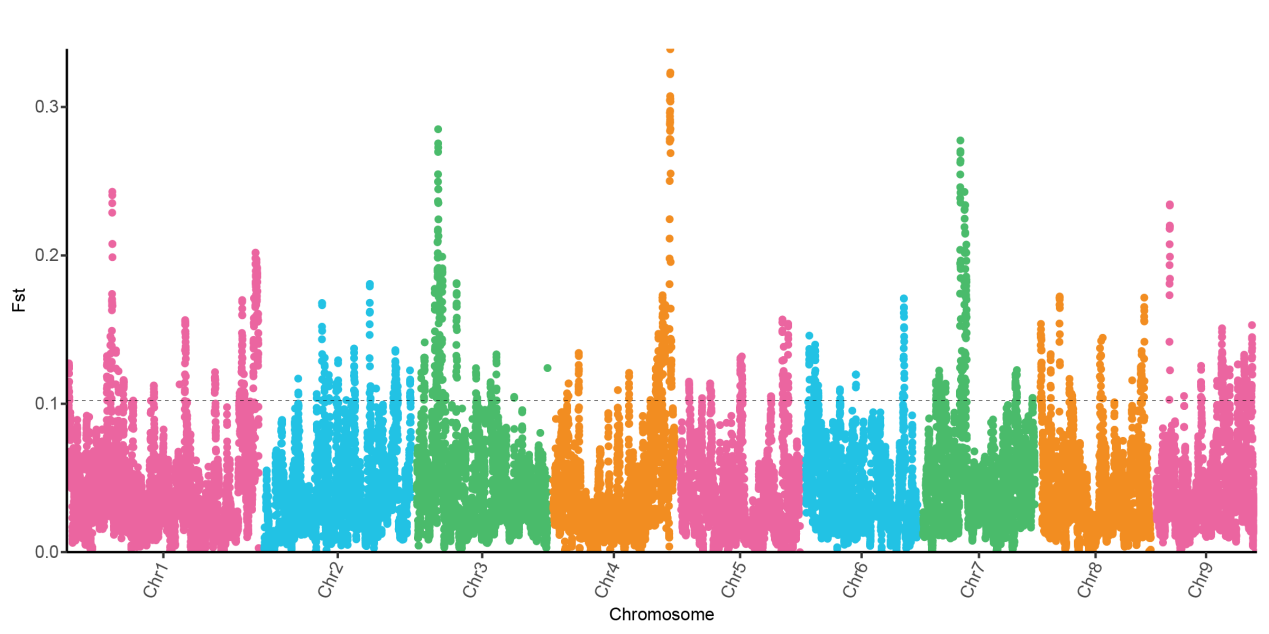

Figure S11. Manhattan plot of genome-wide Fst between landrance and cultivar accessions on the 9 chromosomes. Dotted line, 1% threshold.

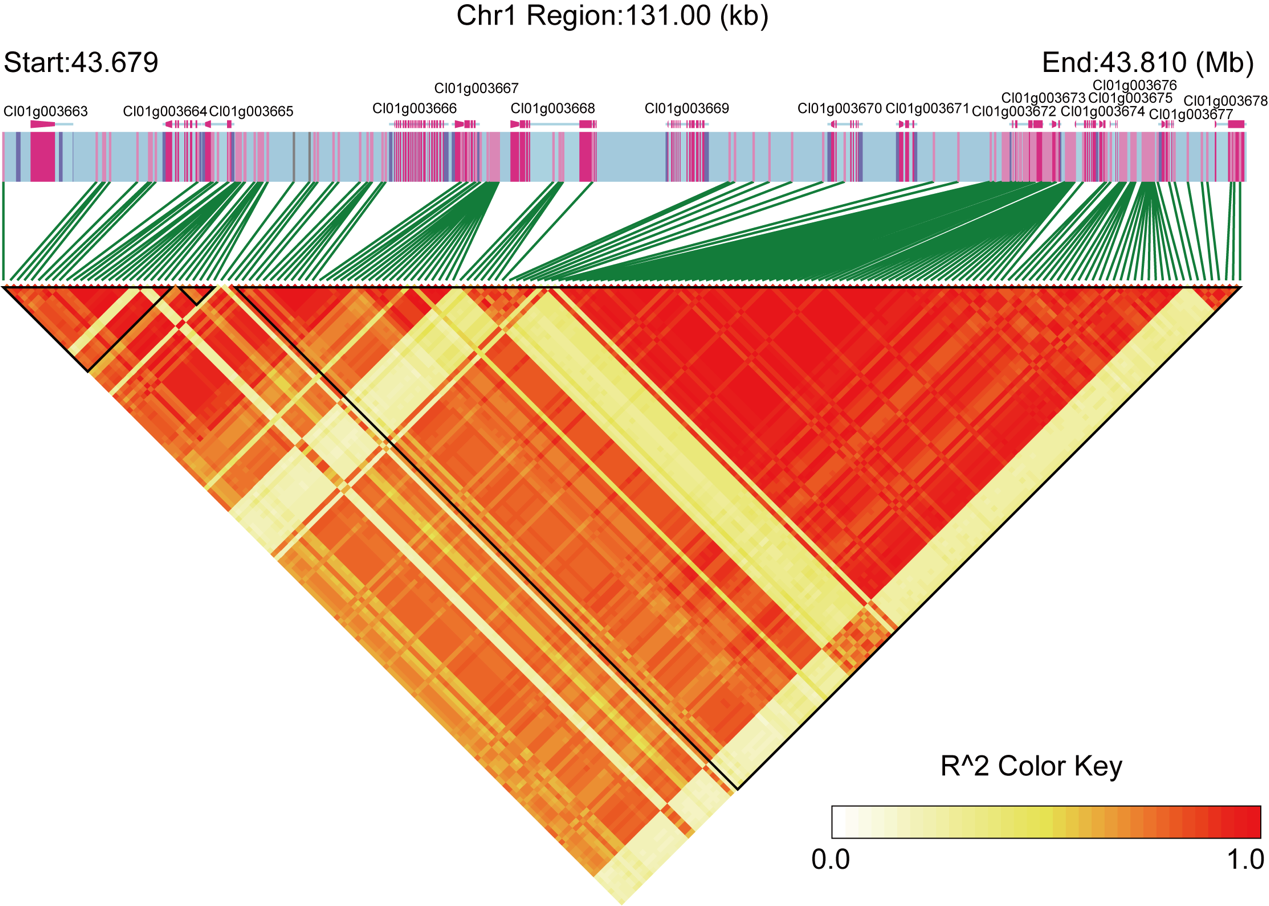

Figure S12. Linkage disequilibrium of all SNPs in the selective region on Chr1 (43700001-43810000) corresponding to the red dots on Figure 4B.

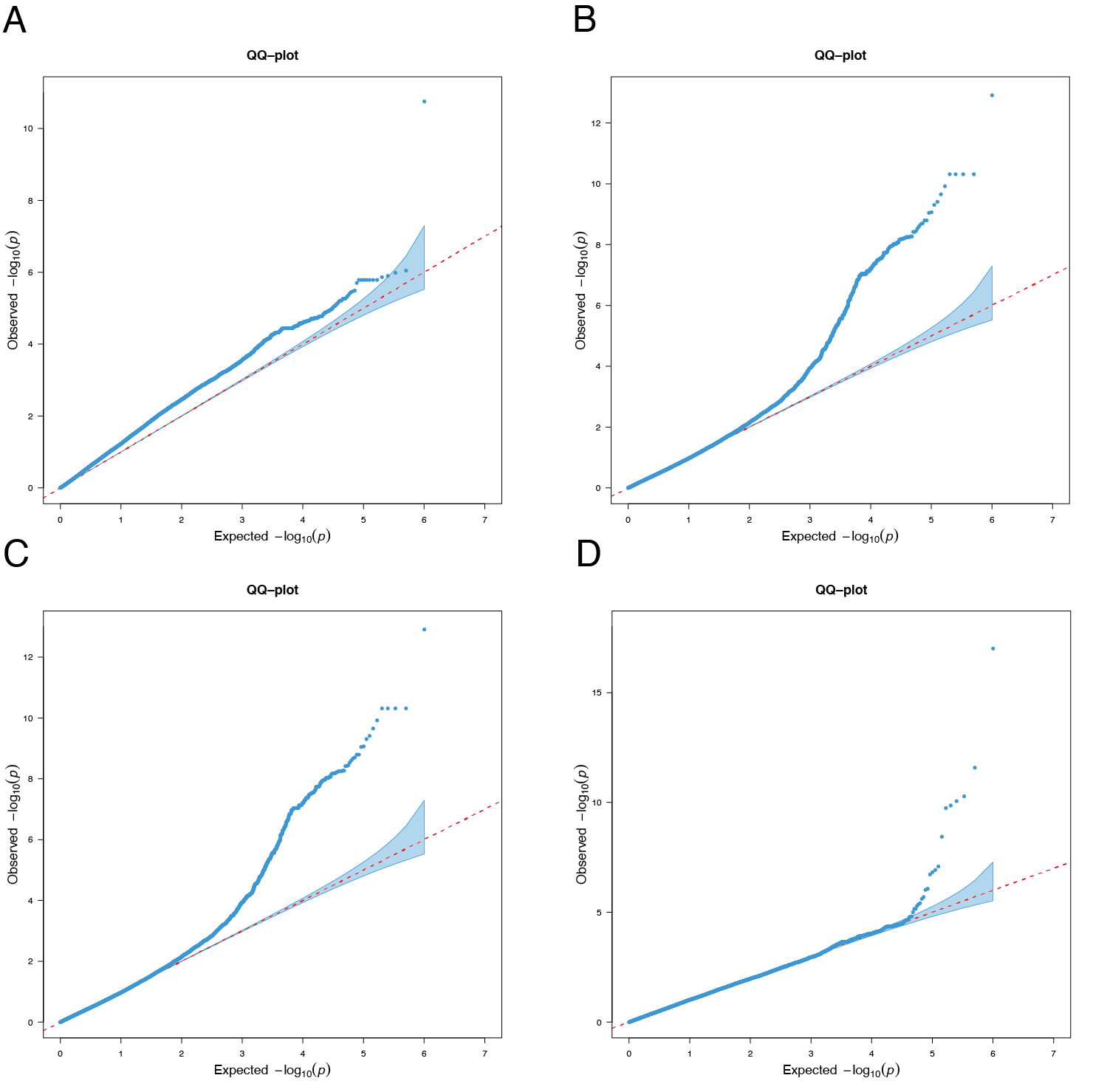

Figure S13. Q-Q plot displaying the GWAS results for total acid content in wampee fruit using: A, Blink; B, CMLM; C, ECMLM; D, FarmCPU models.

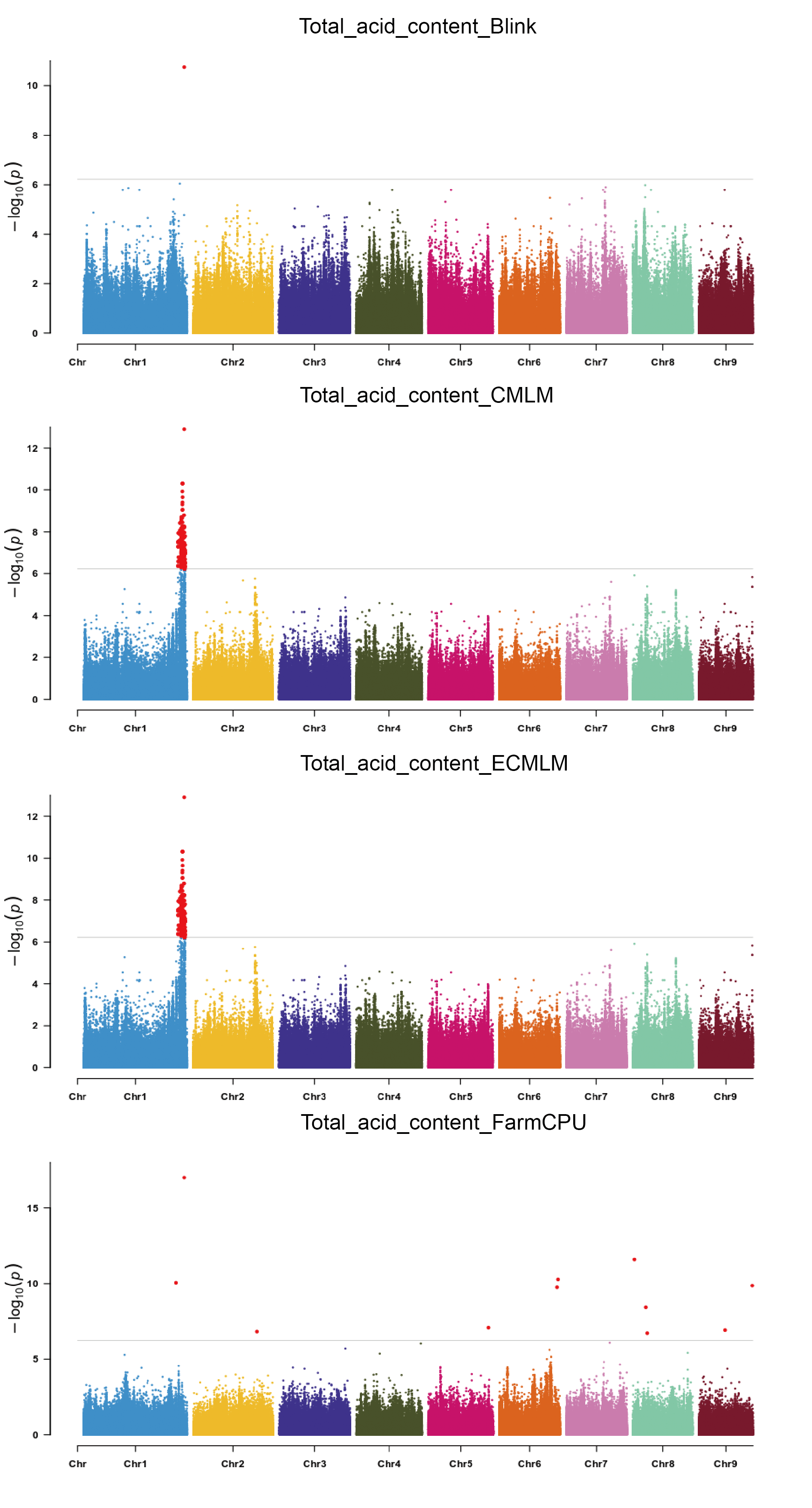

Figure S14. Manhattan plot displaying the GWAS results for total acid content in wampee using FarmCPU, MLMLM, Blink, CMLM, ECMLM models respectively. The threshold for significance is P-value < 6.03E^-7^ (-log10P> 6.22).
